# Competition between parallel sensorimotor learning systems

**DOI:** 10.1101/2020.12.01.406777

**Authors:** Scott T. Albert, Jihoon Jang, Adrian M. Haith, Gonzalo Lerner, Valeria Della-Maggiore, John W. Krakauer, Reza Shadmehr

## Abstract

Sensorimotor adaptation benefits from learning in two parallel systems: one that has access to explicit knowledge, and another that relies on implicit, unconscious correction. However, it is unclear how these systems interact: does enhancing one system’s contributions, for example through instruction, impair the other, or do they learn independently? Here we illustrate that certain contexts can lead to competition between implicit and explicit learning. In some cases, each system is responsive to a task-related visual error. This shared error appears to create competition between these systems, such that when the explicit system increases its response, errors are siphoned away from the implicit system, thus reducing its learning. This model suggests that explicit strategy can mask changes in implicit error sensitivity related to savings and interference. Other contexts suggest that the implicit system can respond to multiple error sources. When these error sources conflict, a second type of competition occurs. Thus, the data show that during sensorimotor adaptation, behavior is shaped by competition between parallel learning systems.

## Introduction

When we reach towards an object, unexpected perturbations to the arm engage multiple corrective systems. Some systems are reactive and respond online to counter the perturbation^1–3^, whereas others are predictive, changing their output to anticipate the perturbation^4–6^. When multiple predictive systems operate together, how do they coordinate their responses to error?

One possibility is that each learning system operates on a separate error source. For example, when people adapt to a visual perturbation and an inertial perturbation simultaneously, the brain engages parallel circuits^7^ that respond to each error separately without interference^8^. In other cases, however, separate corrective systems may respond to a common error. For example, current models suggest that a given sensory error simultaneously engages multiple adaptive systems, each with their own timescale of learning: some fast and others slow^9,10^.

Presence of multiple learning systems in the brain makes it crucial to understand how they are coordinated to seamlessly improve behavior. First, suppose two learning systems are driven by the same error and produce an output that reduces that error (Fig. 1A). In this case, when one system adapts, it reduces the error that is available to drive learning in the other system; thus, these two parallel systems will compete to “consume” a common error. Second, suppose two systems are driven by distinct errors, each producing an output to minimize its own error (Fig. 1B). In this case, when one system adapts to its error, the resulting action could increase the other system’s error, thus producing another type of competition where only one system can minimize its error. These ideas illustrate that a given system’s behavior will depend not only on its own error source, but the error sources that drive parallel learning systems.

**Figure 1.**
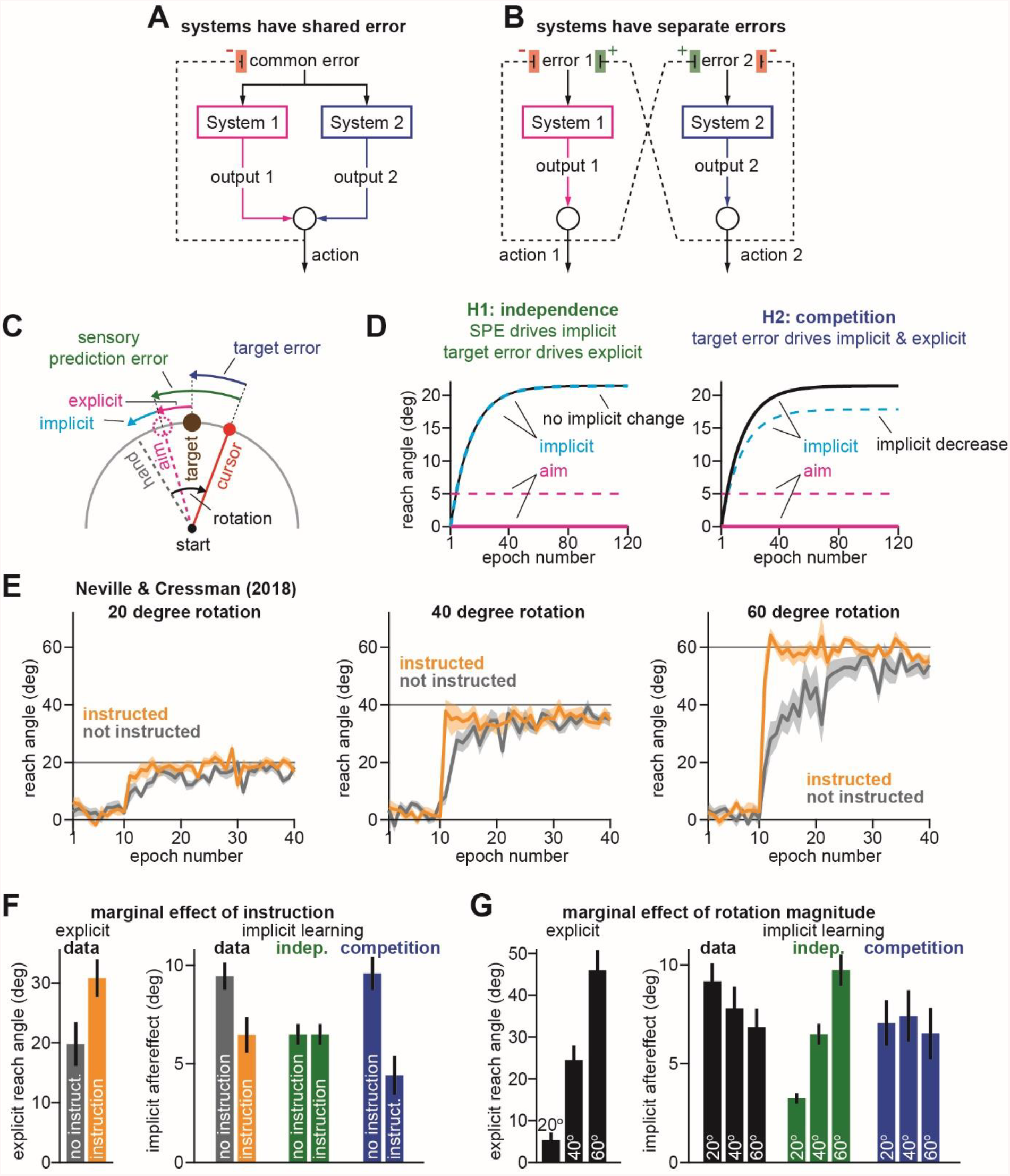
Enhancing explicit strategy suppresses implicit adaptation. **A**. Schematic showing competition between two cooperating parallel systems. Systems 1 and 2 receive the same error and produce outputs to reduce the error. Increases in one system’s output will decrease the error source for the partner system, suppressing its adaptation. **B.** Schematic showing competition between two parallel systems with differing objectives. Systems 1 and 2 receive different errors and produce an output that tends to increase the other system’s error. In this case, when one system is optimized, the other system is prevented from reducing its error. **C.** Schematic of visuomotor rotation. Participants move from S to T. Hand path is composed of explicit (aim) and implicit corrections. Cursor path is perturbed by rotation. We explored two hypotheses: prediction error (H1, aim vs. cursor) vs. target error (H2, target vs. cursor) drives implicit learning. **D.** Prediction error hypothesis predicts that enhancing aiming (dashed magenta) will not change implicit learning (black vs. dashed cyan) according to the independence equation. Target error hypothesis predicts that enhancing aiming (dashed magenta) will decrease implicit adaptation (black vs. dashed cyan). **E.** Data reported by Neville and Cressman^15^. Participants were separated into 1 of 6 groups. Groups differed based on verbal instruction (instructed yellow; non-instructed gray) and rotation magnitude (20° left; 40° middle; 60° right). **F.** The marginal effect of instruction (average across 3 rotation sizes) shown for explicit adaptation at left and implicit learning at right. Learning predicted by the independence equation (green) and competition equation (blue) are shown. Models were fit assuming implicit error sensitivity and retention were identical across all 6 groups. **G.** The marginal effect of perturbation magnitude (average across instruction conditions) shown for explicit adaptation at left and implicit learning at right. Learning predicted by the independence equation (green) and competition equation (blue) are shown. Models were fit as in **F**. Error bars for data show mean ± SEM. Error bars for model predictions refer to mean and standard deviation across 10,000 bootstrapped samples.

Here we consider how these competitive interactions may couple together neural systems that respond to visual errors. Multiple lines of evidence suggest that the brain engages two parallel systems during motor learning: a strategic explicit system that can be guided by instruction^11,12^, as well as an implicit system that adapts without our conscious awareness^12,13^. How might these learning systems interact^14–16^ during sensorimotor adaptation?

The answer depends on their respective error sources. Current models suggest that implicit and explicit systems are differentially engaged by two distinct error sources: a task error^17–19^, and a prediction error^4,12,20^. One theory suggests that the explicit system acts to decrease errors in task performance, while the implicit system acts to reduce errors in predicting sensory outcomes^12,21,22^. However, other models have suggested that both systems are at least partly engaged by errors in task outcome^14,17,23,24^. Here we show that both errors drive implicit learning, but their relative contributions vary across different experiments. Some experiments reveal how learning systems exhibit competition due to a common error source as in Fig. 1A, but in others, they interfere given a conflict between separate errors as in Fig. 1B.

Critically, one’s viewpoint can lead to contrasting interpretations of the same data. Consider the case where implicit and explicit systems share at least one common error source. Suppose some experimental condition facilitates explicit strategy. In this case, increases in explicit strategy will siphon away the error that the implicit system needs to adapt, thus reducing implicit learning without actually changing implicit learning properties.

Changes in implicit learning might occur not solely across two distinct environments, but across two moments in time. For example, when two opposing perturbations are learned in sequence, the rate of learning decreases due to interference^25–27^. On the other hand, when the perturbations are the same, the rate of learning increases due to savings^28–32^. If implicit and explicit systems share an error source, each system’s current response can be shaped not solely by past experience, but also by changes in the other system. This may explain a potential disconnect between studies that have suggested that experience-dependent increases in learning rate are subserved solely by flexible explicit strategies^28,33–36^, and studies that have pointed to concomitant changes in implicit learning systems^17,37,38^.

Here, we mathematically^9,14,24,39,40^ consider the extent to which implicit and explicit systems are engaged by common errors, or separate errors. The hypotheses make diverging predictions, which we then test in various contexts. In some contexts, the data suggest that the two systems are mostly driven by a common error. This shared error produces competition as in Fig. 1A, such that increases^15,16^ or decreases^41,42^ in explicit strategy indirectly exert the opposite effect on implicit learning. This competitive relationship suggests an alternate way that implicit systems may exhibit two hallmarks of learning: savings and interference. However, in other contexts, a single common error cannot explain implicit behavior. In these cases, the data are more consistent with the idea that multiple error sources (e.g., a prediction and a task error) drive comparable levels of implicit learning, leading to competition resembling Fig. 1B.

Together, our results illustrate that changes in behavior during sensorimotor adaptation are shaped by multiple types of competition between parallel learning systems.

## Results

In visuomotor rotation paradigms, participants move a cursor with their hand (Fig. 1C), but experience a perturbation that changes the canonical relationship between hand motion and cursor motion. The perturbation induces adaptation, resulting in a change in reach direction. This adaptation is supported by both implicit and explicit processes^11,12,21,43^; participants can intentionally re-aim their reach angle (Fig. 1C, aim), and also change their reach via implicit recalibration (Fig. 1C, implicit). Together, these two systems determine the hand’s path (Fig. 1C, hand).

Suppose that a rotation *r* alters the cursor’s path (Fig. 1C, cursor). Current models suggest that this perturbation creates two distinct error sources. One error source is created by the deviation between the cursor and the target: a target error^17–19^. Notably, this target error (Fig. 1C, target error) is altered by both implicit (*x_i_*) and explicit (*x_e_*) adaptation:

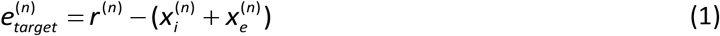

Under normal circumstances, the brain expects that the cursor will move toward the aimed location. This expectation gives rise to a second error: a sensory prediction error (SPE)^4,12,20^. This SPE is created by the deviation between where we aimed our hand (the expected cursor motion) and where we observed the cursor’s actual motion (Fig. 1C, sensory prediction error). Critically, because this error is anchored to our aim location, it is altered solely by changes in the implicit system:

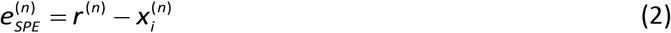

These errors create two different objective functions: (1) maximize success by eliminating target error, and (2) improve our predictions by eliminating SPE. How does the brain’s subconscious learning system respond to these disparate directives? State-space models describe implicit adaptation as a process of learning and forgetting^9,14,24,39,40^:

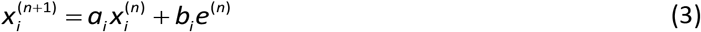

Forgetting is controlled by the retention factor (*a_i_*) which specifies how strongly we retain the adapted state. Learning is controlled by one’s error sensitivity (*b_i_*) which determines the amount we adapt in response to an error – but which error?

To answer this question, consider how Eq. (3) behaves following an extended training period. Like adapted behavior^23,37,44,45^, Eq. (3) approaches an asymptotic limit when the processes of learning and forgetting balance each other (Fig. 1B, implicit). In the extreme case where the implicit system responds solely to target error, total implicit learning is determined by Eqs. (1) and (3):

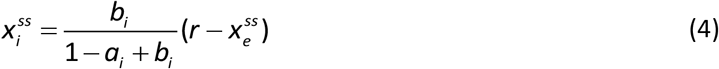

Eq. (4) demonstrates a competition between implicit and explicit systems; the total amount of implicit adaptation 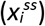 is related to the difference between the perturbation *r* and the total amount of explicit adaptation 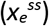.

On the other extreme, when the implicit system responds solely to SPE, total implicit learning is determined by Eqs. (2) and (3):

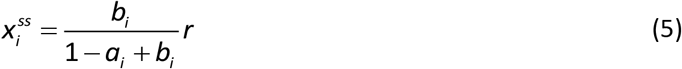

Eq. (5) demonstrates an independence between implicit and explicit systems; the total amount of implicit adaptation depends solely on the rotation’s magnitude, not one’s explicit strategy.

In summary, the competition (Eq. (4)) and independence (Eq. (5)) equations make predictions that can answer a critical question: which errors drive implicit adaptation? If implicit learning is predominantly driven by SPE, the implicit system will depend only on the perturbation’s magnitude according to the independence equation (Eq. (5)). On the other hand, if implicit learning is predominantly driven by target error, the implicit system will compete with explicit strategies according to the competition equation (Eq. (4)). Here, we investigate these predictions across several experimental paradigms and explore their limitations in describing the behavior of the implicit learning system.

### Enhancement in explicit strategy reduces the amount of implicit adaptation

Suppose that in one condition, participants adapt to a visual rotation with some fixed explicit strategy (Fig. 1D, aim, solid magenta line). But in a second condition, the participant is coached about the perturbation^16,46^, enhancing their explicit strategy (Fig. 1D, aim, dashed magenta line). If the implicit system learns only from SPE (Eq. (5)), then changes in explicit strategy will have no impact on implicit learning (Fig. 1D, H1, compare solid black and dashed blue implicit lines). On the other hand, if the implicit system learns only from target error, it competes with the explicit system (Eq. (4)). Coaching explicit strategy suppresses implicit learning (Fig. 1D, H2, compare dashed blue and solid black implicit lines).

To test this prediction, we considered an experiment performed by Neville and Cressman^15^. Participants were exposed to either a 20°, 40°, or 60° visuomotor rotation (Fig. 1E), and separated into instructed and non-instructed conditions. Non-instructed groups (Fig. 1E, gray) adapted without any initial instruction regarding the perturbation. Instructed participants were briefed about the upcoming rotation and how they should compensate to hit the target (Fig. 1E, yellow). This instruction sharply increased the rate of adaptation over that of the non-instructed group (Fig. 1E, compare yellow and gray curves).

To determine how instruction accelerated adaptation, participants were asked to reach with and without explicit strategy (Fig. S1). The marginal effects of instruction (average across rotation magnitudes) and perturbation magnitude (average over instruction conditions) are shown in Figs. 1F and 1G respectively. Unsurprisingly, instructed participants learned faster due to an enhancement in explicit reaiming, which increased by approximately 10° across each rotation magnitude (Fig. 1F, explicit).

Curiously, while instruction enhanced explicit learning, it appeared to impair implicit adaptation, decreasing the total implicit aftereffect (Fig. 1F, implicit learning, data). Even more puzzling, whereas contributions of the explicit system increased with rotation magnitude (Fig. 1G, explicit), implicit learning did not, as one might intuitively expect (Fig. 1G, implicit learning, data).

To interpret the implicit response to awareness and perturbation magnitude, we fit both the competition (Eq. (4)) and independence equations (Eq. (5)) to the behavior across all groups, under the assumption that the implicit system’s sensitivity to error and retention (*b_i_* and *a_i_*) were identical across all rotation sizes, and across the instructed and non-instructed conditions.

The independence and competition models made contrasting predictions (see individual predictions in Figs. S1B&C). Because SPE does not depend on explicit aiming, Eq. (5) incorrectly predicted the same level of implicit learning irrespective of explicit awareness (Fig. 1F, implicit learning, indep.). Furthermore, because implicit adaptation is driven solely by the rotation magnitude in the independent model, Eq. (5) also incorrectly predicted that implicit learning should increase with rotation size (Fig. 1G, implicit learning, indep.).

The opposite was true of the competition model. Eq. (4) correctly predicted less implicit learning in instructed participants who used greater explicit strategy (Fig. 1F, implicit learning, competition). Remarkably, the competition model also predicted that the implicit aftereffect should remain similar across rotation magnitudes (Fig. 1G, implicit learning, competition). How was this possible? Critically, the competition equation suggests that the driving force for implicit learning is not solely the rotation, but the difference between the rotation and explicit strategy. Therefore, because the total amount of explicit reaiming increased as the rotation magnitude increased (Fig. 1G, explicit), their difference remained roughly constant across all perturbation sizes (Fig. S1D). Thus, Eq. (4) predicted similar implicit aftereffects irrespective of rotation size.

In summary, when explicit learning is enhanced through instruction, implicit learning is impaired. As perturbation magnitude increases, contributions of explicit learning increases, but not the contributions of implicit learning. These observations are consistent with the competition model (Eq. (4)), suggesting that the implicit and explicit systems are primarily driven by a common target error.

### Suppression of explicit learning increases the amount of implicit adaptation

The competition equation predicts that enhancing explicit strategy should decrease implicit learning (Fig. 1). What should happen when explicit learning is suppressed? Suppose participants adapt with an explicit strategy (Fig. 2B, aim, solid magenta line), but this strategy is then suppressed (Fig. 2B, aim, dashed magenta line). Because SPE learning does not depend on explicit strategy, Eq. (5) predicts no change in implicit learning (Fig. 2B, H1, left, compare solid black and dashed blue implicit lines) (Eq. (5)). However, because target errors do depend on explicit strategy, Eq. (4) predicts that suppressing explicit aiming will increase implicit learning (Fig. 2B, H2, right, compare dashed blue and solid black implicit lines).

**Figure 2.**
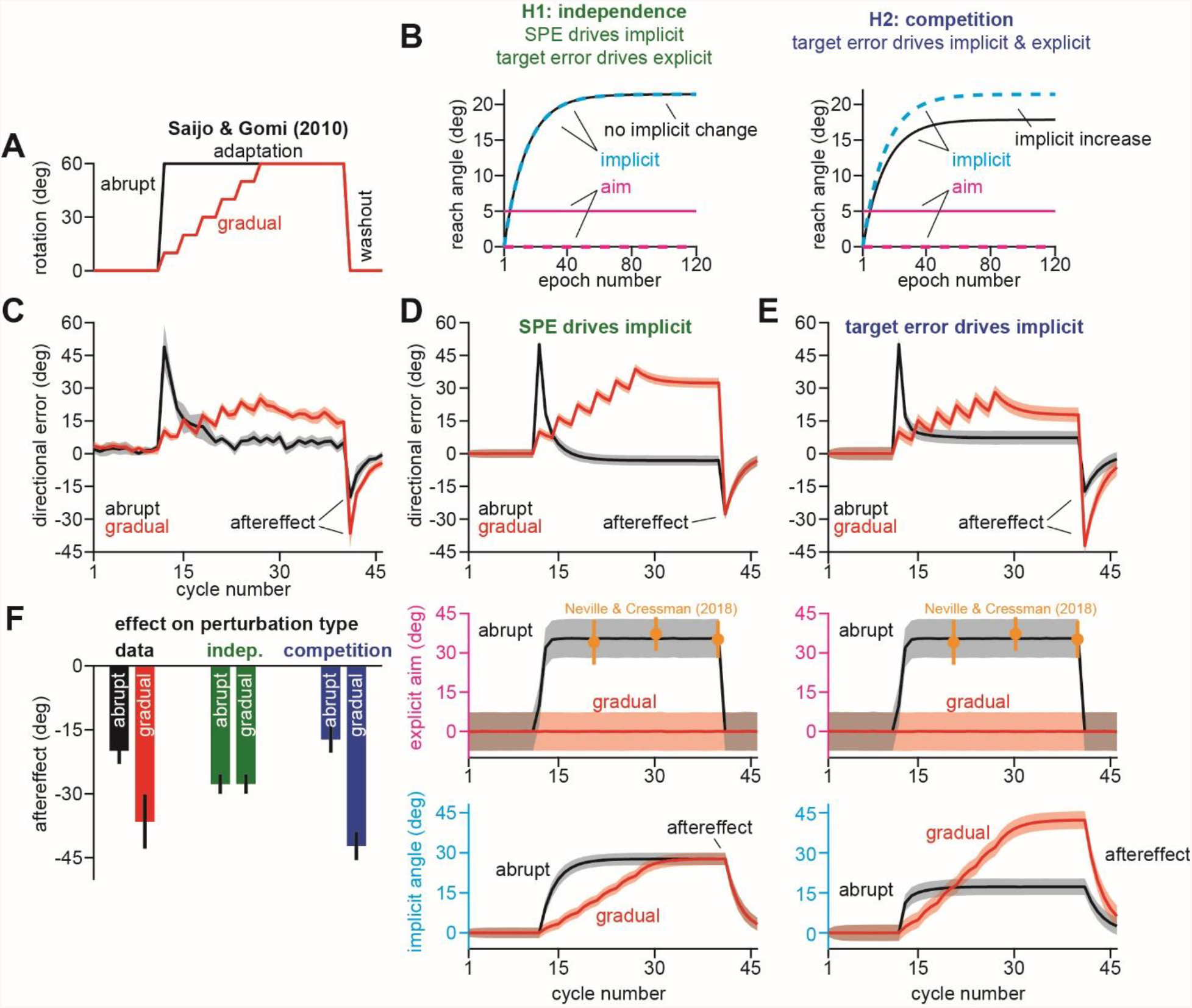
Suppressing explicit strategy increases the total amount of implicit adaptation. Data reported from Saijo and Gomi^42^. **A.** Participants adapted to either an abrupt or gradual 60° rotation followed by a washout period. **B.** We explored two hypotheses: prediction error (H1, aim vs. cursor) vs. target error (H2, target vs. cursor) drives implicit learning. Prediction error hypothesis predicts that suppressing aiming (dashed magenta) through gradual perturbation onset will not change implicit learning (black vs. dashed cyan). Target error hypothesis predicts that suppressing aiming (dashed magenta) will increase implicit adaptation (black vs. dashed cyan). **C.** Directional error during adaptation. Note that while the abrupt group exhibited greater adaptation during the rotation, they also showed a smaller aftereffect suggesting less implicit adaptation. **D.** We simulated a state-space model where the implicit system learned from SPE. The model parameters were selected to best fit the data in **C**. In the middle row, hypothetical abrupt explicit strategy was simulated based on data reported by Neville and Cressman^15^ (yellow points). The gradual explicit strategy was assumed to be zero because participants were less aware. At bottom, we show implicit learning predicted by an SPE error source. Note the identical saturation levels. **E.** Same as in **D**, but for implicit adaptation based on target error. Note greater implicit learning in gradual condition at the bottom row. Models in **D** and **E** were fit assuming that implicit error sensitivity and retention are identical across abrupt and gradual conditions. **F.** Here we show the implicit aftereffect on the first washout cycle (12 total trials). Model predictions for SPE learning (indep.) and target error learning (competition) are shown. Data show mean ± SEM across participants. Error bars for model are mean and standard deviation across 20,000 bootstrapped samples.

One way to suppress explicit learning is to make participants unaware by introducing the perturbation gradually. In an earlier experiment, Saijo and Gomi (2010)^42^ exposed participants to either an abrupt (Fig. 2A, abrupt) or gradual (Fig. 2A, gradual) perturbation. The abrupt perturbation was immediately set to 60°, but the gradual perturbation reached this magnitude over time.

Participants in the abrupt condition adapted rapidly to the perturbation, greatly decreasing their target error to about 5° over about 10 perturbation cycles (Fig. 2C, abrupt). Participants in the gradual group, experienced small target errors throughout training, but adapted less by the end of the rotation period, exhibiting a terminal error nearly 3 times greater than the abrupt condition (Fig. 2C, gradual).

At this point, the perturbation was abruptly removed, revealing large aftereffects in each group. However, even though participants in the gradual group had adapted less completely to the rotation, they paradoxically exhibited larger aftereffects (Fig. 2F, data), which remained elevated throughout the entire washout period (Fig. 2C, aftereffect). If these aftereffects reveal the total amount of implicit adaptation, given that strategies are rapidly disengaged when the perturbation is removed^34^ (Fig. S2), how could more complete adaptation in the abrupt group lead to less implicit adaptation?

To investigate this phenomenon, we considered how implicit and explicit systems might behave according to the independence (Eq. (5)) and competition (Eq. (4)) frameworks. To simulate these models, we estimated the explicit strategies in each group. Neville and Cressman^15^ had measured the explicit response to a 60° rotation, demonstrating that participants re-aimed their hand approximately 35° consistently over the adaptation period (see yellow points in Figs. 2D&E, explicit aim). This estimate agreed well with the data; participants in the abrupt condition adapted 55°, and exhibited an aftereffect of approximately 20° (Fig. 2F, data, abrupt), suggesting about 35° of re-aiming. In the gradual group, we assumed that little to no re-aiming occurred. This also seemed consistent with the data; participants in the gradual group adapted approximately 40°, and exhibited an aftereffect of approximately 38° (Fig. 2F, data, gradual) suggesting <5° of re-aiming. Using these estimates, we constructed hypothetical explicit learning timecourses, as shown in Figs. 2D&E, explicit aim).

We next used the state-space model to simulate the implicit learning timecourse, in cases where the implicit system learned solely due to SPE (Fig. 2D, implicit angle) or solely due to target error (Fig. 2E, implicit angle), under the assumption that participants in both the abrupt and gradual groups had the same implicit error sensitivity (*b_i_*) and retention factor (*a_i_*). The parameter sets that yielded the closest match to the measured behavior (Fig. 2C) are shown in Figs. 2D&E (directional error). In both cases, the models predicted abrupt and gradual learning timecourses that resembled the data.

However, the implicit states predicted by SPE learning and target error learning possessed a critical difference. According to Eq. (4), the target error model predicted that the total extent of implicit learning would be suppressed by explicit strategy in the abrupt condition, yielding a smaller aftereffect (Fig. 2E, implicit angle). However, according to Eq. (5), the SPE model predicted that implicit learning should reach the same level, yielding identical aftereffects (Fig. 2D, implicit angle).

In summary, the differences in aftereffects across the abrupt and gradual conditions (Fig. 2F, data) were accurately predicted by the competition model (Fig. 2F, competition), but not the independence model (Fig. 2F, indep.). Suppressing the explicit strategy revealed competition between implicit and explicit systems which suggested that the implicit system predominantly responded to target error.

### Subject-to-subject correlations reveal competition between implicit and explicit systems

Data in Figs. 1 and 2 suggested that the implicit system was altered by competition with explicit strategy. Is this competition observed at the level of individual participants? In other words, the competition model would predict that participants who use larger strategies will naturally exhibit less implicit adaptation.

To investigate this possibility, we considered earlier work where Fernandez-Ruiz and colleages^41^ exposed participants to a 60° rotation (Fig. 3A). The large rotation appeared to induce substantial variation in strategic re-aiming. Consider for example Subjects A and B (Figs. 3B&C). Upon rotation onset, Subject A rapidly reduced their directional error (Fig. 3B, Subject A) and exhibited two characteristics that suggested the use of large explicit re-aiming angles: (1) their reach angle varied greatly from one cycle to the next^14,44,47^ and (2) their movement preparation time (Fig. 3C, Subject A) greatly increased upon onset of the perturbation^18,28,37,47^. On the other hand, Subject B reduced directional errors slowly and consistently (Fig. 3B, Subject B), with little to no increase in movement preparation time (Fig. 3C, Subject B). Thus, Subjects A and B appeared to engage explicit strategies to differing extents. How did differences in their explicit strategy impact implicit learning?

**Figure 3.**
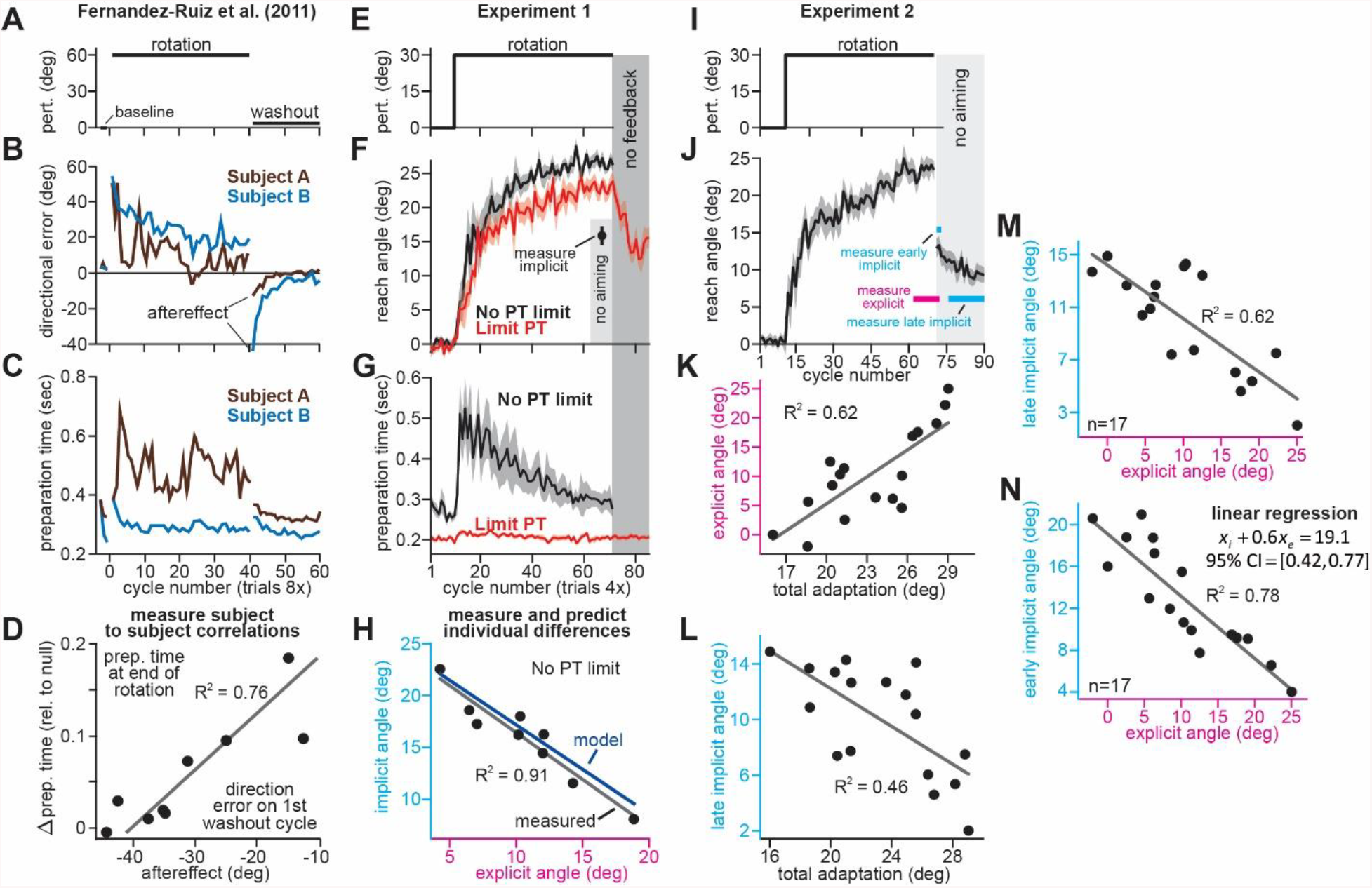
Strategy suppresses implicit learning across individual participants. **A-D.** Data are shown from Fernandez-Ruiz et al.^41^. Participants were exposed to a 60° visuomotor rotation followed by a washout period. Paradigm shown in **A**. Two learning curves for individual participants shown in **B**. Preparation time (latency between reach onset and target presentation) shown in **C**. In **D**, participants with greater increases in preparation time (relative to baseline) showed smaller aftereffects, suggesting less implicit adaptation. **E-F.** In Experiment 1, participants adapted to a 30° visuomotor rotation. The paradigm is shown in **E**. Participants in the No PT limit group had no constraint placed on their movement preparation time. Participants in the Limit PT group had to execute movements with restricted preparation time. Learning curves for each group shown in **F**. Note that Limit PT adaptation ended with a no feedback period where memory retention was measured. Note that No PT limit adaptation ended with a cycle of exclusion trials where participants were instructed to reach straight to the target without re-aiming and without any feedback (no aiming, measure implicit). Movement preparation time for each group is shown in **G**. In **H**, we show the total implicit and explicit adaptation in each participant in the No PT limit condition. Implicit learning measured during the terminal no aiming probe. Explicit learning represents difference between total adaptation (last 10 rotation cycles) and implicit probe. The black line shows a linear regression. The blue line shows the theoretical relationship predicted by the competition equation which assumes implicit system adapts to target error. The parameters for this model prediction (implicit error sensitivity and retention) were measured in the Limit PT group. **I-N.** In Experiment 2, participants performed a similar experiment remotely using a personal computer. The paradigm is shown in **I**. The learning curve is shown in **J**. Implicit learning was measured at the end of adaptation over a 20-cycle period where participants were instructed to reach straight to the target without aiming and without feedback (no aiming seen in **I** and **J**). We measured explicit adaptation as difference between total adaptation and reach angle on first no aiming cycle (**J**, measure explicit). We measured early implicit aftereffect as reach angle on first no aiming cycle (**J**, measure early implicit). We measured late implicit aftereffect as mean reach angle over last 15 no aiming cycles (**J**, measure late implicit). In **K** we show how explicit adaptation varies with total adaptation. In **L** we show how late implicit aftereffect varies with total adaptation. In **M** we show how explicit adaptation varies with late implicit aftereffect. In **N** we show how explicit adaptation varies with early implicit aftereffect. Points in **I**-**N** show individual participants. Lines indicate linear regressions. Error bars show mean ± SEM across participants.

When the perturbation was removed, reaction time returned to baseline levels (Fig. 3C), revealing each participant’s aftereffect (Fig. 3B, aftereffect). Paradoxically, though Subject A adapted more completely to the rotation during the adaptation period, they exhibited a far smaller aftereffect (Fig. 3B). A possible explanation is that because Subject A used greater explicit strategy during adaptation, their implicit system adapted less due to competition, producing a smaller aftereffect. Indeed, although participants who increased their preparation time exhibited smaller reach errors (Fig. S3), engaging explicit strategies appeared to inhibit their implicit system, as revealed by a decrease in the aftereffect during the washout period (Fig. 3D; *ρ*=0.87, p<0.01).

The competition model (Eq. (4)) provides a way to quantify these subject-to-subject correlations. The left-most term in this equation is a learning gain that varies between 0 and 1, which depends on implicit learning properties: retention (*a_i_*) and error sensitivity (*b_i_*). Thus, the competition equation predicts that implicit and explicit learning will negatively co-vary according to a line whose slope and bias are determined by the properties of the implicit learning system (*a_i_* and *b_i_*). To test the model’s accuracy, we exposed participants to a 30° visuomotor rotation (Fig. 3E) under two conditions (Experiment 1). In one group, we strictly limited preparation time to inhibit time-consuming explicit strategies^41,47^ (Fig. 3F, Limit PT). In the other group, we imposed no preparation time constraints (Fig. 3F, No PT limit). Our goal was to measure *a_i_* and *b_i_* in the Limit PT group which putatively relied on implicit learning, and use these values to predict the implicit-explicit relationship across No PT limit participants.

As expected, PT Limit participants dramatically reduced their reach latencies throughout the adaptation period, whereas No PT limit participants exhibited a sharp increase in movement preparation time after perturbation onset (Fig. 3G), indicating explicit re-aiming^18,28,37,41,47^. Consistent with suppression of explicit strategy, learning proceeded more slowly and was less complete with the PT Limit (Fig. 3F; two-sample t-test on last 10 adaptation epochs: t(30)=2.14, p=0.041, d=0.77).

Next, we empirically measured the putative implicit retention factor (*a_i_*) and error sensitivity (*b_i_*) associated with the PT Limit learning curve. We measured the retention factor during a terminal “no feedback” period (Fig. 3F, dark gray, no feedback) and error sensitivity (*b_i_*) during the adaptation period (see Methods). Together, this retention factor (*a_i_*=0.943) and error sensitivity (*b_i_*=0.35), produced a specific form of Eq. (4), namely, *x_i_* = 0.86 (30 – *x_e_*), which we could use to predict how implicit and explicit learning should vary across participants in the No PT limit group (Fig. 3H, blue line).

To measure No PT limit implicit and explicit learning we instructed participants to move their hand through the target without any re-aiming at the end of the adaptation period (Fig. 3F, no aiming). The precipitous change in reach angle revealed the terminal amounts of implicit and explicit adaptation (post-instruction reveals implicit; total drop reveals explicit). To verify the accuracy of this explicit measure, we asked participants to verbally report their re-aiming angles (see Methods). Participants that demonstrated greater explicit strategy indeed reported larger re-aiming angles at the end of adaptation (Fig. S4A, *ρ*=0.709) and also appeared to require greater movement preparation time (Fig. S4B, *ρ*=0.708).

How did subject-to-subject variations in implicit and explicit learning compare to the model’s prediction? We observed a striking correspondence between the No PT limit implicit-explicit relationship (Fig. 1H, black dot for each participant; *ρ*=−0.95) and that predicted by the competition model (Fig. 3H, blue). The slope and intercept predicted by Eq. (4) (−0.86 and 25.74°, respectively) differed from the measured linear regression (Fig. 1H, black line, R^2^=0.91; slope = −0.9 with 95% CI [−1.16, −0.65] and intercept = 25.46° with 95% CI [22.54°, 28.38°]) by only about 5% and 1%, respectively.

Lastly, we tested two alternate explanations that could also explain the observed correlations between implicit and explicit learning. First, explicit (total adaptation minus no aiming probe) and implicit (no aiming probe) learning measures inherently share variance which could lead to spurious correlation. Second, in the event that participants exhibit nearly identical learning asymptotes, say approximately 26° in our experiment, these implicit and explicit learning measures could be trivially constrained to lie along the regression line: *x_i_* + *x_e_* ≈ C, where C = 26°.

To test these possibilities, we conducted a control experiment (Experiment 2). Participants adapted to a 30° rotation again (Fig. 3I), but this time, we measured implicit adaptation using the no-aiming instruction over an extended 20-cycle period (Fig. 3J, no aiming). We calculated early (first no-aiming cycle; Fig. 3J, measure early implicit) and late (last 15 no-aiming cycles; Fig. 3J, measure late implicit) implicit learning measures. As in Fig. 3H, we calculated total explicit strategy as the difference between total adaptation and the first no-aiming cycle (Fig. 3J, measure explicit).

Critically, our explicit measure and late implicit measure were now properly decoupled, as they depended on separate cycles. Remarkably, late implicit learning exhibited patterns that matched the group-level interventions observed by Neville and Cressman^15^ (Fig. 1) and Saijo and Gomi^42^ (Fig. 2). Namely, participants that compensated most for the perturbation utilized large explicit strategies (Fig. 3K; *ρ*=0.79, p<0.001). But enhancements in overall learning came at the cost of reductions in implicit adaptation (Fig. 3L; *ρ*=−0.68, p=0.003), due to a competition between implicit and explicit learning (Fig. 3M*, ρ*=−0.79, p<0.001).

Secondly, we considered the relationship between explicit strategy and early implicit learning, and again observed a strong negative linear relationship (Fig. 3L, *ρ*=−0.79): *x_i_* + 0.6*x_e_* = 19.1. Notably, the explicit regression coefficient’s (0.6) 95% CI, [0.42,0.77] did not contain 1. Equivalently, this indicates that there was substantial variation in asymptotic learning across participants (range 16-29°), ruling out the trivial possibility that *x_i_* + *x_e_* = C, described above. To the contrary, participants who showed greater explicit learning had better overall compensation for the perturbation, but had less implicit learning.

In summary, consistent with the idea that the two learning systems share a common error, we found that when a subject’s performance depends more on the contributions of the explicit system, their implicit system learns less.

### Competition predicts increases in both implicit and explicit error sensitivity during savings

When participants are exposed to the same perturbation twice, they adapt more quickly the second time. This phenomenon is known as savings and is a hallmark of sensorimotor adaptation^9,48,49^. Multiple studies have attributed this process solely to changes in explicit strategy^28,33,34,36,50^.

For example, in an earlier work^28^, we trained participants (n=14) to reach to one of two targets, coincident with an audio tone (Fig. 4A). By shifting the displayed target approximately 300 ms prior to tone onset on a minority of trials (20%), we forced participants to execute movements with limited preparation time (Low preparation time; Fig. 4A, middle). On trials in which subjects had high preparation time, i.e. trials without a target switch (Fig. 4B, left), adaptation exhibited savings; the rate of learning increased across exposures (Fig. 4B, right, High PT; Wilcoxon signed rank, p=0.0085, Cohen’s d=0.683). Learning differences were most pronounced on the first 40 trials after perturbation onset (Fig. 4C, left; Fig. 4C, right, paired t-test, p=0.0044, Cohen’s d=0.920).

**Figure 4.**
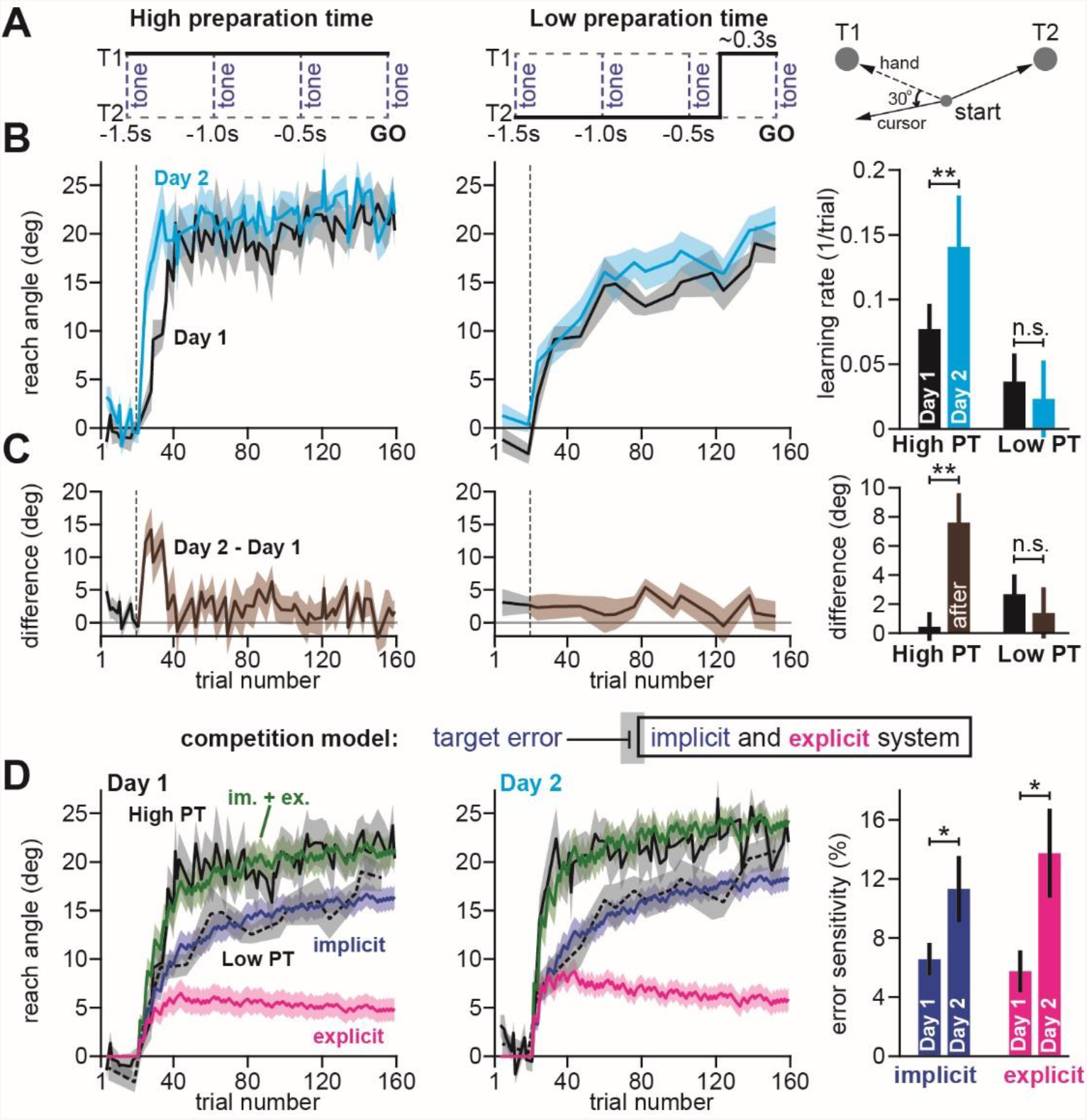
Model predicts increase in implicit error sensitivity without any change in implicit learning rate. **A.** Haith and colleagues^28^ instructed participants to reach to Targets T1 and T2 (right). Participants were exposed to a 30° visuomotor rotation at Target T1 only. Participants reached to the target coincident with a tone. Four tones were played with a 500 ms inter-tone-interval. On most trials (80%) the same target was displayed during all four tones (left, High preparation time or High PT). On some trials (20%) the target switched approximately 300 ms prior to the fourth tone (middle, Low preparation time or Low PT). **B.** On Day 1, participants adapted to a 30° visuomotor rotation (Block 1, black) followed by a washout period. On Day 2, participants again experienced a 30° rotation (Block 2, blue). At left, we show the reach angle expressed on High PT trials during Blocks 1 and 2. Dashed vertical line shows perturbation onset. At middle, we show the same but for Low PT trials. At right, we show learning rate on High and Low PT trials, during each block. **C.** As an alternative to the rate measure shown at right in **B**, we calculated the difference between reach angle on Blocks 1 and 2. At left and middle, we show the learning curve differences for High and Low PT trials, respectively. At right, we show difference in learning curves before (black) and after (brown) the perturbation. **D.** We fit a state space model to the learning curves in Blocks 1 and 2 assuming that target errors drove implicit adaptation. Low PT trials captured the implicit system (blue). High PT trials captured the sum implicit and explicit system (green). Explicit trace (magenta) is the difference between the High and Low PT predictions. At right, we show error sensitivities predicted by the model. Error bars show mean ± SEM, except for the learning rate in **B** which displays the median. Paired t-tests are used in **C** and **D**. Wilcoxon signed rank test is used in **B**. Statistics: n.s. means no significant difference, *p<0.05, **p<0.01.

To test for changes in implicit learning, we focused on short PT trials where explicit strategy is suppressed^41,47^. Unlike the High PT trials, adaptation expressed on short PT trials was similar during the two exposures (Fig. 4B, middle); we found no difference in the rate of short PT learning (Fig. 4B, right, Wilcoxon signed rank, p=0.903). Similarly, the difference in learning curves for exposures 1 and 2 (Fig. 4C, middle) did not show any change after perturbation onset (Fig. 4C, right, Low PT, paired t-test, p=0.624).

These results suggested that savings relied solely on a time-consuming explicit strategy. Does this mean that implicit learning was completely unaltered by prior exposure to the perturbation? The answer depends on which errors drive implicit adaptation.

In the competition model, implicit learning is driven by target errors (Eq. (1)) that are also shared with the explicit system. We fit this model to the behavior of each participant under the assumption that the reach angle on low preparation time trials revealed the implicit state of adaptation, and the reach angle on high preparation time trials revealed the sum of the implicit and explicit states of adaptation. The model generated implicit (Fig. 4D, left and middle, blue) and explicit (Fig. 4D, left and middle, magenta) states that tracked the behavior well in high PT trials (Fig. 4D, left and middle, solid black line) as well as low PT trials (Fig. 4D, left and middle, dashed black line).

Unsurprisingly, given that High PT trials exhibited savings but Low PT trials did not, the model predicted that explicit error sensitivity increased across exposures, thus leading to an increased rate of adaptation (Fig. 4D, right, explicit; paired t-test, p=0.016, Cohen’s d=0.738). However, the model unmasked a surprising possibility; even though the implicit system showed no increase in learning rate on Low PT trials (Figs. 4B&C, right), the model still indicated that the implicit system had increased its error sensitivity across exposures (Fig. 4D, right, implicit, paired t-test, p=0.023, Cohen’s d=0.686).

In contrast, when we fit the same data assuming that implicit adaptation was driven by SPE rather than target error (Eq. (2), learning depends on rotation but not explicit strategy), the model (not shown in Fig. 4) predicted that only explicit (paired t-test, p=0.026, Cohen’s d=0.673) but not implicit (paired t-test, p=0.099) error sensitivity had increased.

In summary, when we reanalyzed our earlier data, Eqs. (4) and (5) suggested that the same data could be interpreted in two different ways. If we assumed that implicit learning is independent of explicit strategy (independence equation), then only explicit strategy contributed to savings. This is in fact what we had concluded in our original report. However, if we assumed that the implicit and explicit systems learned from the same error (competition equation), then both implicit and explicit systems contributed to savings. How can we determine which interpretation is more parsimonious with measured behavior?

### Competition with explicit strategy can alter measurement of implicit learning

Suppose you arrive at your family dinner, but on this occasion are feeling particularly famished. Yet after the meal, you are surprised to find that you ate the same amount as last week despite feeling hungrier. Does this mean your hunger level was actually the same? No, not necessarily; because you are sharing the meal with others, changes in their consumption rates alter the food available to you. So, eating the same amount could mean that your sister sitting next to you was also hungrier than usual, taking more than their normal share, and thus leaving less for you.

The competition equation (Eq. (4)) presents an analogous scenario, except here the “family” in question is the implicit and explicit adaptive states, and the “food” that is available for consumption is error. The competition model provides the insight that when the explicit system learns faster than before (Fig. 4D, Day 2 vs. Day 1), it leaves less error to drive implicit learning. However, despite this reduced error for the implicit system, performance on Low PT trials on Day 2 was comparable to Day 1 (Fig. 4B, right). Thus, error sensitivity of the implicit system must also have increased from Day 1 to Day 2.

To understand how our ability to detect changes in implicit adaptation can be altered by explicit strategy we constructed a competition map (Fig. 5A). Imagine that we want to compare behavior across two timepoints or conditions. Fig. 5A shows how change in implicit error sensitivity (x-axis) and explicit error sensitivity (y-axis) both contribute to measured implicit aftereffects (denoted by map colors), based on the competition equation (Eq. (4)). The left region of the map (cooler colors) denotes combinations of implicit and explicit changes that decrease implicit adaptation. The right region of the map (hotter colors) denotes combinations that increase implicit adaptation. The middle black region represents combinations that manifest as a perceived invariance in implicit adaptation (<5% absolute change in implicit adaptation).

**Figure 5.**
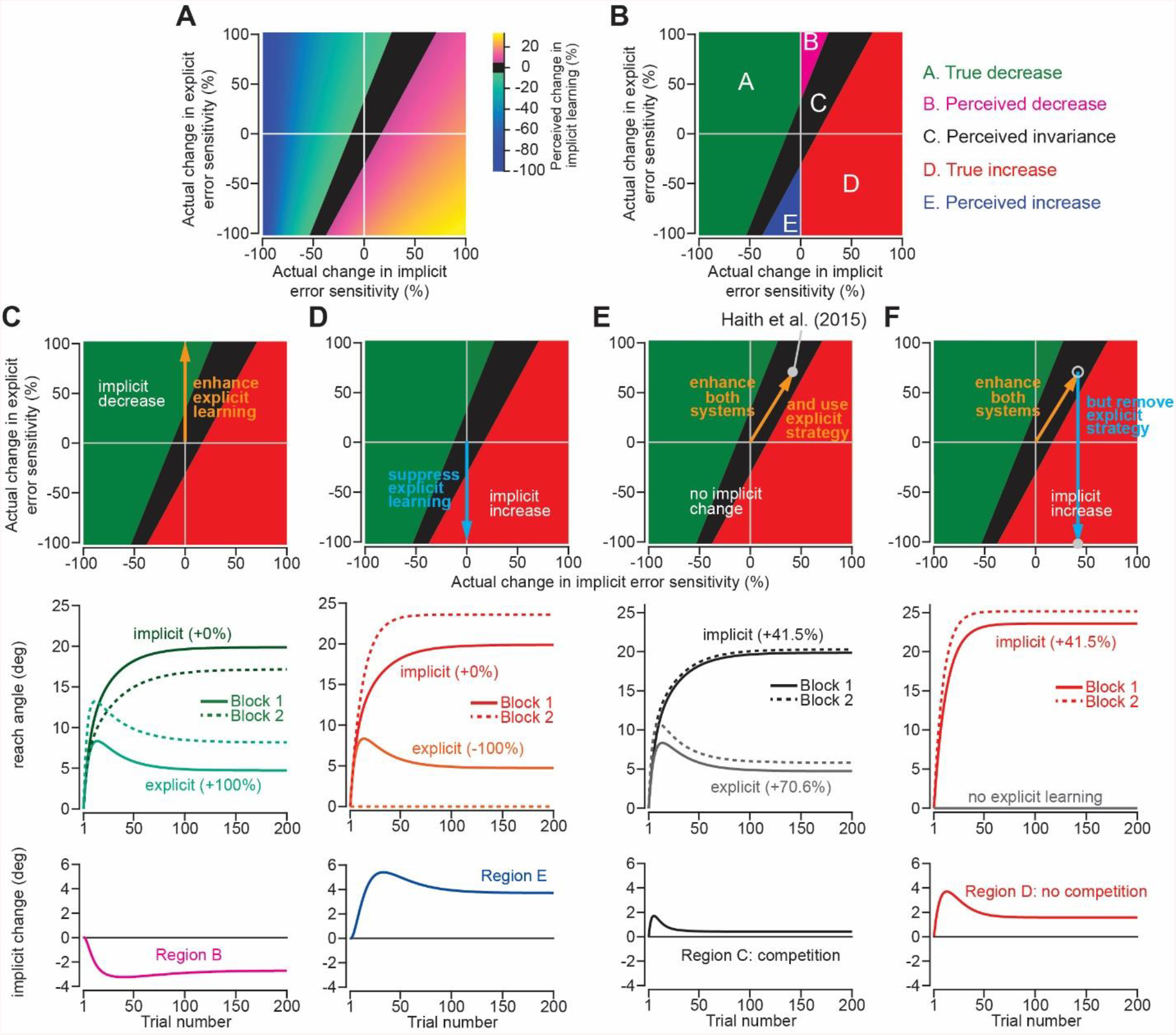
Changes in implicit adaptation depend on both implicit and explicit error sensitivity. **A.** Here we depict the competition map. The x-axis shows change in implicit error sensitivity between reference and test conditions. The y-axis shows change in explicit error sensitivity. Colors indicate the percent change in implicit adaptation (measured at steady-state) from the reference to test conditions. Black region denotes an absolute change less than 5%. The map was constructed with Eq. (8). **B.** The map can be described in terms of 5 different regions. In Region A (true increase), implicit error sensitivity and total implicit adaption both increase in test condition. Region D is same, but for decreases in error sensitivity and total adaptation. In Region B (perceived decrease) implicit adaption decreases though its error sensitivity is higher or same. In Region E (perceived increase), implicit adaptation increases though its error sensitivity is lower or same. Region C shows a perceived invariance where implicit adaptation changes less than 5%. **C.** Top: effect of suppressing explicit learning. Middle: implicit and explicit learning shown in Blocks 1 and 2, where explicit error sensitivity increases 100%. Bottom: implicit learning change (Block 1 to 2). **D.** Top: effect of enhancing explicit learning. Middle: implicit and explicit learning shown in Blocks 1 and 2, where only difference is 100% increase in explicit error sensitivity. Bottom: change in implicit learning (Block 1 to 2). **E.** Top: model simulation for Haith et al.^28^. Middle: implicit and explicit learning during Blocks 1 and 2 where implicit error sensitivity increases by 41.5% and explicit error sensitivity increases by 70.6%. Bottom: negligible change in implicit learning (Block 1 to 2). **F.** Same as in **E** except here explicit strategy is suppressed during Blocks 1 and 2.

Practically, this map defines several distinct regions (Fig. 5B). In Region A, there is a “true decrease” in implicit adaptation; that is, implicit error sensitivity decreases between Timepoints 1 and 2 as does the total amount of implicit learning. Region D is similar, but for simultaneous increases in implicit error sensitivity and total implicit learning (“true increase”).

The other regions describe more surprising situations. In Region B, there is only a “perceived decrease” in implicit learning; that is, implicit learning decreases, even though the implicit error sensitivity has actually increased or remained the same. In Region E, there is only a “perceived increase” in implicit learning; implicit learning increases, even though its error sensitivity decreased or remained the same.

Indeed, we have already explored these phenomena in Figs. 1 and 2. In Fig. 1, enhancing explicit strategy decreased implicit learning without changing any implicit learning properties. The scenario is equivalent to moving up the y-axis of the map (Fig. 5C, top). The same implicit system will decrease its output (Fig. 5C, bottom) when normal levels of explicit strategy are increased (Fig. 5C, middle). On the other hand, suppressing explicit strategy by gradually changing the perturbation appeared to increase implicit learning without changing any implicit learning properties (Fig. 2). This scenario is equivalent to moving down the y-axis of the map (Fig. 5D, top). The same implicit system will increase its output (Fig. 5D, bottom) when normal levels of explicit strategy are then suppressed (Fig. 5D, middle).

Now, let us consider the savings task in Fig. 4. The target error-driven (Eq. (1)) state space model predicted (Fig. 3D) that explicit error sensitivity increased by approximately 70.6% during the second exposure, whereas the implicit system’s error sensitivity increased by approximately 41.5% (Fig. 5E, middle). These changes in implicit and explicit adaptation describe a single point in the competition map, denoted by the gray circle in Fig. 5E (top). This experiment occupies Region C, which indicates that despite the 41.5% increase in implicit error sensitivity, the total amount of implicit learning will increase by less than 5% (Fig. 5E, bottom). In other words, the competition equation suggests the possibility that savings could have occurred in the implicit system but was hidden by a dramatic increase in explicit strategy.

To test this prediction, we can suppress explicit adaptation, thus eliminating competition (Fig. 5F, middle). Such an intervention would move our experiment from Region C to Region D (Fig. 5F, top) where we will observe greater change in the implicit process (Fig. 5D, bottom). Thus, we performed a new experiment to test this prediction.

### Savings in implicit learning is unmasked by suppression of explicit strategy

The key prediction is that removal of explicit strategy will unmask savings in implicit learning (Fig. 5F). We exposed participants (Experiment 3) to two 30° rotations, separated by an intervening period of washout. To suppress explicit strategy, we forced participants to move under strict reaction time constraints on every trial. As a result, participants reached to each of the four targets with a latency of approximately 200 ms (Fig. 6B, top), nearly 100 ms sooner than the Low PT condition used in our earlier experiment^28^ (Fig. 6A). When reaction time was limited on all trials, the learning rate during the second exposure (Fig. 6B, middle) exhibited a marked increase (Fig. 6C, no comp.; Wilcoxon signed rank, p=0.014, Cohen’s d=0.637). This enhancement in learning developed immediately after perturbation onset (Fig. 6B, bottom; Fig. 6C, bottom, no comp., paired t-test, p=0.008, Cohen’s d=1.06).

**Figure 6.**
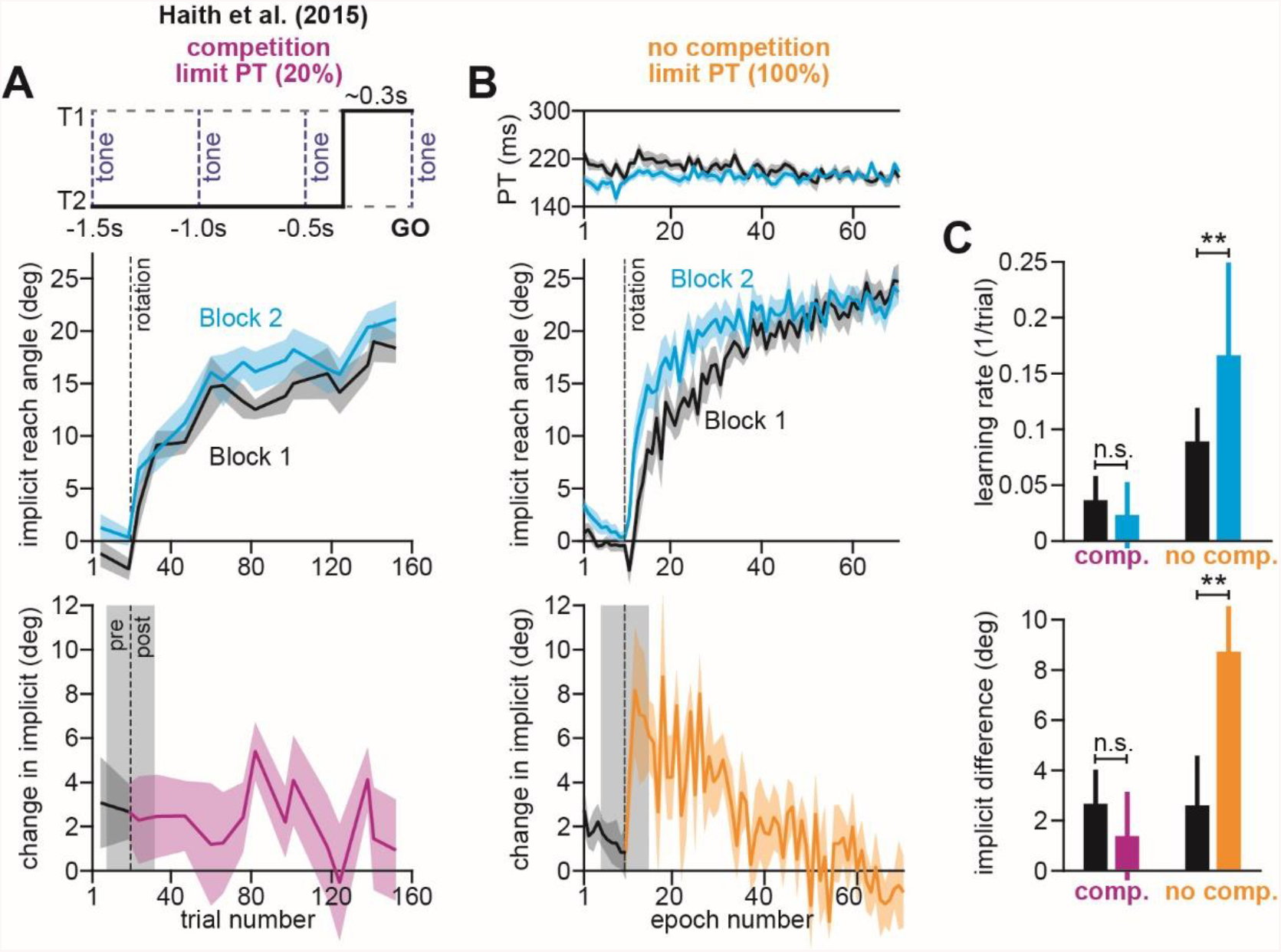
Removing explicit strategy reveals savings in implicit adaptation. **A**. Top: Low preparation time (Low PT) trials in Haith and colleagues^28^ used to isolate implicit learning. Middle: learning during Low PT in Blocks 1 and 2. Bottom: difference in Low PT learning between Blocks 1 and 2. **B.** Similar to **A**, but here (Experiment 3) explicit learning was suppressed on every trial, as opposed to only 20% of trials. To suppress explicit strategy, we restricted reaction time on every trial. The reaction time during Blocks 1 and 2 is shown at top. At middle, we show how participants adapted to the rotation under constrained reaction time. At bottom, we show the difference between the learning curves in Blocks 1 and 2. **C.** Here we measured savings in Haith et al. (20% of trials had reaction time limit) and Experiment 3 (100% of trials had reaction time limit). At top, we quantify savings by fitting an exponential curve to each learning curve. Bars show the rate parameter associated with the exponential. At bottom, we quantify savings by comparing how Blocks 1 and 2 differed before perturbation onset (black), and after perturbation onset (purple and yellow). Error bars show mean ± SEM, except for the learning rate at the top of **C** which shows the median. Paired t-tests are used at the bottom of **C**. Wilcoxon signed rank tests are used at the top of **C**. Statistics: n.s. means no significant difference, **p<0.01.

In summary, when explicit learning was suppressed, Low PT behavior exhibited savings (Fig. 6B). But when explicit strategies remained active, Low PT behavior did not exhibit any change in learning rate (Fig. 6A). One possible explanation for these observations is that an implicit system expressible at Low PT exhibits savings, but this can be masked by competition with explicit strategy.

### Impairments in implicit learning lead to anterograde interference

When two opposing perturbations (say *A* and *B*) are experienced in sequence, exposure to perturbation *A* decreases the rate of learning in *B* (anterograde interference). Like savings^29,32,48,49^, we recently suggested that impaired learning in *B* is caused by a change in error sensitivity^26^. Might this change in error sensitivity depend on the implicit learning system?

We exposed two groups of participants to opposing visuomotor rotations of 30° and −30° in sequence (Experiment 4). In one group, the perturbations were separated by a 5-minute break (Fig. 7A). In a second group, the break was 24 hours in duration (Fig. 7B). We suppressed explicit strategies by strictly limiting reaction time. Under these constraints, participants executed movements at latencies slightly greater than 200 ms (Figs. 7A&B, middle, blue). These reaction times were approximately 50% lower than those observed when no reaction time constraints were imposed on participants, as in Albert & Lerner et al.^26^ (Figs. 7A&B, middle, green).

**Figure 7.**
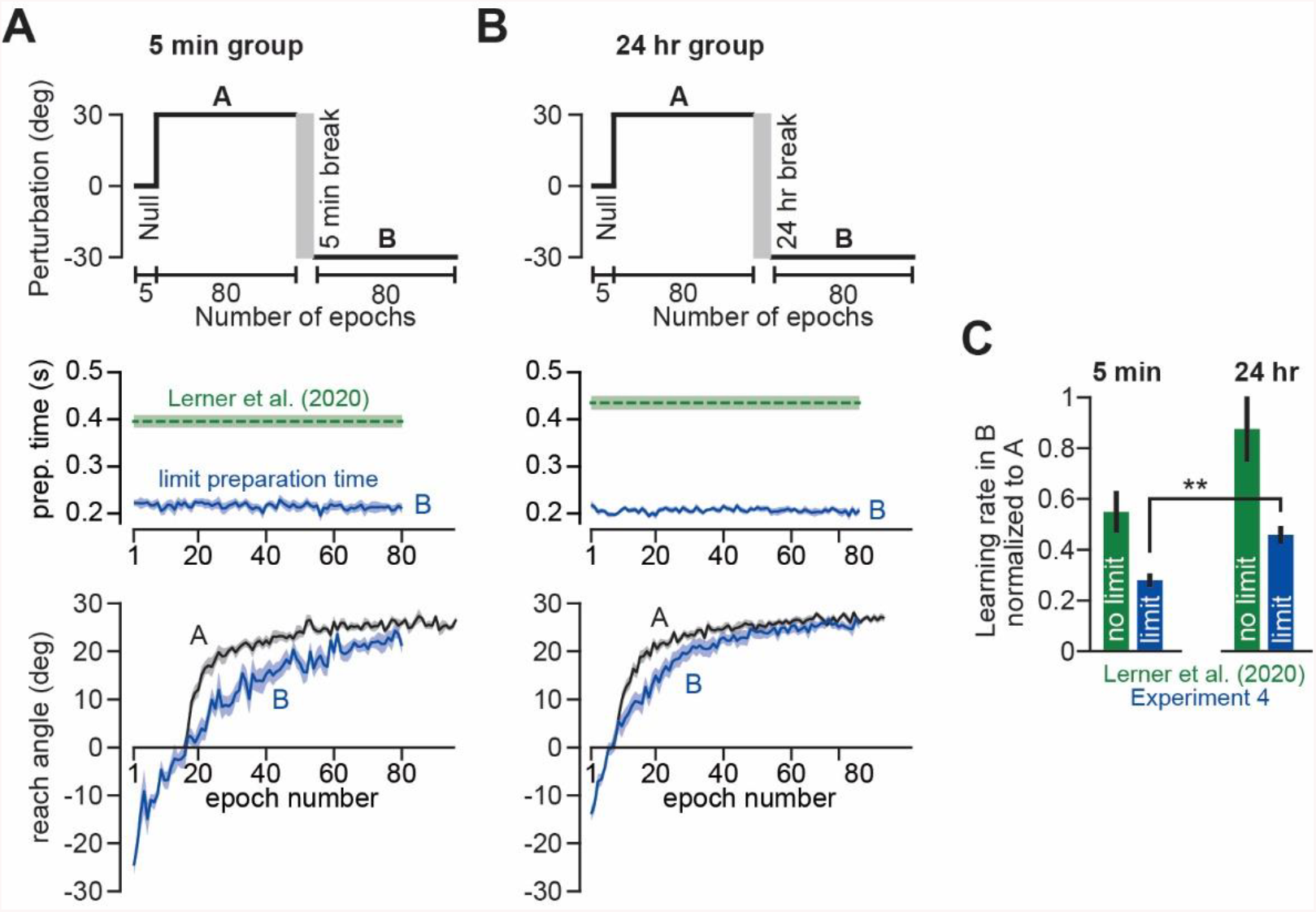
Removing explicit strategy reveals anterograde interference in implicit adaptation. **A.** Top: participants were adapted to a 30° rotation (A). Following a 5-minute break, participants were then exposed to a −30° rotation (B). This A-B paradigm was similar to that of Lerner & Albert et al.^26^. Middle: to isolate implicit adaptation, we imposed strict reaction time constraints on every trial. Under these constraints, reaction time (blue) was reduced by approximately 50% over that observed in the self-paced condition (green) studied by Lerner & Albert et al.^26^ Bottom: learning curves during A and B in Experiment 4; under reaction time constraints, the interference paradigm produced a strong impairment in the rate of implicit adaptation. To compare learning during A and B, B period learning was reflected across y-axis. Furthermore, the curves were temporally aligned such that an exponential fit to the A period and exponential fit to the B period intersected when the reach angle crossed 0°. This alignment visually highlights differences in the learning rate during the A and B periods. **B.** Here we show the same analysis as in **A** but when exposures A and B were separated by 24 hours. **C.** To measure the amount of anterograde interference on the implicit learning system, we fit an exponential to the A and B period behavior. Here we show the B period exponential rate parameter divided by the A period rate parameter (values less than 1 indicate a slowing of adaptation). At left we show the results for the 5-minute group. At right we show the results for the 24-hr group. In green we show data from Lerner & Albert et al.^26^ where reaction time was unrestricted (no limit). In blue we show our new dataset (Experiment 4) where reaction time was limited to isolate implicit learning. A two-sample t-test was used to test for differences in the implicit impairment at 5 minutes and 24 hours. Error bars show mean ± SEM. Statistics: **p<0.01.

We found that implicit adaptation during the second rotation period was significantly impaired after a 5-minute break (Fig. 7A, bottom). The rate of implicit learning decreased by approximately 75% (Fig. 7C, 5min, limit). Passage of time partially improved this deficit (Fig. 7B, bottom). When the rotations were separated by a 24 hr break, implicit learning rate was impaired by only 55% (Fig. 7C, 24 hr, limit).

Thus, we can conclude that suppression of explicit strategy revealed an anterograde deficit in implicit learning that did not completely resolve after 24 hours, perhaps even stronger than that observed when no reaction time constraints are imposed^26^ (Fig. 7C, Lerner et al. (2020), no limit; see Discussion).

### The implicit system may adapt to multiple target errors at the same time

The idea that a single shared error drives both implicit and explicit learning is quite surprising. After all, in earlier work by Mazzoni and Krakauer^12^, it appeared that implicit learning was driven by outcome-independent prediction errors (Eq. (2)) that were unaltered by explicit strategy. Yet, in Figs. 1–7, implicit learning clearly depended, at least in part, on target error, and exhibited clear interactions with explicit strategy. How does one reconcile the current results with the results of Mazzoni and Krakauer?

To explore this question, we revisited these earlier experiments. In Mazzoni and Krakauer, we tested two sets of participants. In the no-strategy group, participants adapted to a standard 45° rotation (Fig. 8A, blue, no-strategy, adaptation) followed by washout (Fig. 8A, blue, no-strategy, washout). In a second group, participants made two initial movements with the rotation (Fig. 8A, red, strategy, 2 movements no instruction). Then we told participants to aim towards a neighboring target (45° away) which entirely compensated for the rotation. Unlike the experiments described in Figs. 1–7, in which only the primary target was visible, in Mazzoni and Krakauer both the primary target and the aiming target were always visible. Participants immediately adopted the aiming strategy, bringing error with respect to the primary target to zero (Fig. 8A, red, strategy, instruction). Surprisingly, after eliminating this error, their movement angles gradually drifted beyond the primary target, overcompensating for the rotation. These involuntary changes implicated an implicit process.

**Figure 8.**
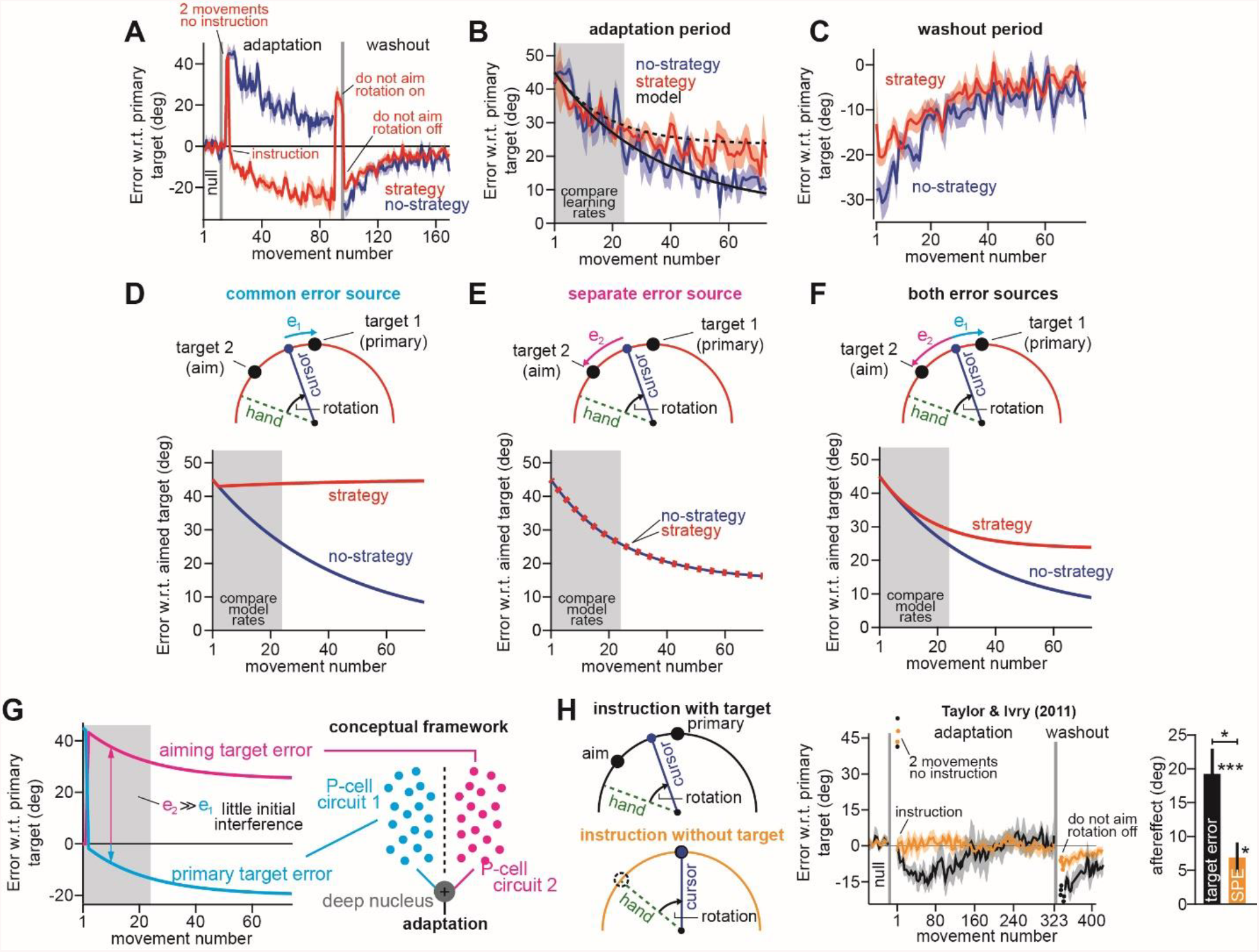
Two visual targets create two implicit error sources. **A.** Figure shows data reported in Mazzoni and Krakauer^12^. Blue shows error between primary target and cursor during adaptation and washout. Red shows the same, but in a strategy group that was instructed to aim to a neighboring target (instruction) to eliminate target errors, once participants experienced two large errors (2 cycles no instruction). **B.** Here we show the error between the cursor and the aimed target during the adaptation period. These curves are the same as in **A** except we use the aimed target rather than primary target, so as to better compare learning curves across groups. **C.** The washout period reported in **A**. Here error is relative to primary target, though in this case aimed and primary targets are the same. **D.** Here we modeled behavior when implicit learning adapts to primary target errors. The primary target error is shown in *e_1_* at top. Note that no-strategy learning resembles data. However, strategy learning exhibits no drift because the implicit system has zero error. Note here that the primary target error of 0° is a 45° aimed target error in the strategy group. **E.** Similar to **D**, except here the implicit system adapts to errors between the cursor and aimed target. This error is schematized in *e_2_* at top. Note that this model predicts identical learning in strategy and no-strategy groups. **F.** In this model, the strategy group adapts to both the primary target error and the aimed target error (*e_1_* and *e_2_* at top). The no-strategy group adapts only to the primary target error. Learning parameters are identical across groups. **G.** At left, we show how aiming target and primary target errors evolve in the strategy group in **F**. At right, we imagine a potential neural substrate for implicit learning. The primary target error and aiming target error engage two different sub-populations of Purkinje cells in the cerebellar cortex. These two implicit learning modules combine at the deep nucleus. **H.** Figure shows data reported in Taylor and Ivry^21^. Participants performed a task similar to **A**. Before adaptation, participants were taught how to re-aim their reach angles. In the “instruction with target” group, participants re-aimed during adaptation with the aide of neighboring aiming targets (top-left). In the “instruction without target” group, participants re-aimed during adaptation without any aiming targets, solely based on the remembered instruction from the baseline period. The middle shows learning curves. In both groups, the first 2 movements were uninstructed, resulting in large errors (2 movements no instruction). Note in the “instruction with target” group, there is an implicit drift as in **A**, but participants eventually reverse this by changing explicit strategy. There is no drift in the “instruction without target” group. At right, we show the implicit aftereffect measured by telling participants not to aim (first no feedback, no aiming cycle post-adaptation). Greater implicit adaptation resulted from physical target. Error bars show mean ± SEM. Statistics: *p<0.05, ***p<0.001.

When we compared the rate of learning with and without strategy, we found that it was not different over the initial exposure to the perturbation (Fig. 8B, gray, compare learning rates; compare mean adaptation over first 24 movements, two-sample t-test, p=0.223). This suggested that implicit adaptation was unaltered by the abrupt change in explicit strategy, and equally importantly, was not driven by error between the cursor and target (Eq. (1)), but rather by a sensory prediction error (Eq. (2)).

However, there remained an unsolved puzzle. While the initial rates of adaptation were the same irrespective of strategy, adaptation diverged later in learning (Fig. 8B, compare strategy and no-strategy curves after the initial gray region; two-sample t-test, p<0.005), with the no-strategy group achieving greater implicit learning (see aftereffect in Fig. 8C; two-sample t-test, p<0.005). Might these late differences have been caused by participants in the strategy group abandoning the explicit strategy as it led to larger and larger errors? This possibility seemed unlikely. When we asked participants to stop re-aiming (Fig. 8A, do not aim rotation on) their movement angle changed by 47.8° (difference between 3 movements before and 3 movements after instruction), indicating that they had continued to maintain the instructed explicit re-aiming strategy near 45°.

We wondered if interactions between implicit and explicit learning could help solve these puzzles. First, we considered the competition model that best described the experiments in Figs. 1–7. In this model, the implicit system is driven exclusively by error with respect to the primary target (Eq. (1)), which is shared with explicit strategy (Fig. 8D, top, *e1*). While this model predicted learning in the standard no-strategy condition, it failed to account for the drift observed when participants were given an explicit strategy (Fig. 8D, no learning in strategy group). This was not surprising. If implicit learning is driven by the primary target’s error, it will not adapt in the strategy group because participants explicitly reduce target error to zero at the start of adaptation (note that −45° in Fig. 8D actually means a 0° primary target error).

We next considered the possibility that implicit learning was driven exclusively by error with respect to the aimed target (target 2, Fig. 8E, top, *e_2_*), as we concluded in our original study^12^. While this model correctly predicted implicit learning in both the no-strategy and strategy conditions, it could not account for any differences in learning that emerged later during the adaptation period (Fig. 8E, bottom).

Finally, we noted that participants in the strategy group were given two contrasting goals. One goal was to aim for the secondary target, whereas the other goal was to move the cursor through the primary target (both targets were always visible). Therefore, we wondered if participants in the strategy group learned from two distinct errors: cursor with respect to target 1, and cursor with respect to target 2 (Fig. 8F, top). In contrast, participants in the no-strategy group attended solely to the primary target, and thus learned only from the error between the cursor and target 1. Thus, we imagined that implicit learning in the strategy group was driven by the two different kinds of target error:

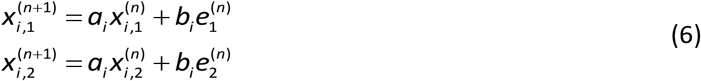

These two modules then combined to determine the total amount of implicit learning (i.e., *x_i_* = *x_i,1_* + *x_i,2_*).

Remarkably, when we applied the dual target error model (Eq. (6)) to the strategy group, and the single target error model (Eqs. (1) & (3)) to the no-strategy group, the same implicit learning parameters (*a_i_* and *b_i_*) closely tracked the observed group behaviors (black model in Fig. 8B). These models correctly predicted that initial learning would be similar across the strategy and no-strategy groups (compare curves in gray region in Fig. 8F bottom), but would diverge later during adaptation. How was this possible?

In Fig. 8G (left), we show how the errors with respect to the primary target and the aiming target evolve as a function of time for the dual target model. Due to the instructed strategy, primary target error is reduced to zero at the start of adaptation (see Fig. 8G, original target error curve). Therefore, early in learning, the implicit system is driven predominantly only by one error source in both the strategy and no-strategy groups, leading to similar adaptation rates. However, as the error with respect to the aimed target decreases, error with respect to the primary target increases but in the opposite direction (Fig. 8G; see schematic in Fig. 8F for intuition). Therefore, the primary target error opposes further adaptation to the aiming target error. This counteracting force causes implicit adaptation to saturate prematurely. Hence, participants in the no-strategy group, who do not experience this error conflict, adapt more.

This re-analysis suggests that when people move a cursor to one visual target (Objective 1), while aiming at another visual target (Objective 2), each target appears to contribute a separate implicit error source. When these two error sources conflict with one another, the implicit learning system can exhibit an unintuitive attenuation in the total amount of adaptation. Thus, while explicit strategies can suppress implicit learning via competition (Figs. 1–7), a different type of suppression can occur when parallel implicit learning systems attempt to solve two conflicting objectives, as in Fig. 1B.

### The persistence of sensory prediction error, in the absence of target error

Our re-analysis in Figs. 8A-G, suggested that when participants use a second target to aim their reach, this additional landmark creates a second implicit error source. To what extent does this error depend on the target’s physical presence in the workspace? Taylor & Ivry^21^ tested this idea, repeating the instruction paradigm used by Mazzoni and Krakauer, though with nearly 4 times the number of adaptation trials (Fig. 8H, instruction with target, black). Interestingly, while the reach angle exhibited the same implicit drift described by Mazzoni and Krakauer, with many more trials participants eventually counteracted this drift by modifying their explicit strategies, bringing their target error back to zero (Fig. 8H, black). At the end of adaptation, participants exhibited large implicit aftereffects after being instructed to no longer aim (Fig. 8H, right, aftereffect; t(9)=5.16, p<0.001, Cohen’s d=1.63).

However, in a second experiment, participants were taught how to re-aim their reach angles during an initial baseline period, but during adaptation itself, they were not provided with physical aiming targets (Fig. 8H, instruction without target). Thus, in this case, only an SPE could drive implicit adaptation towards the aimed location. Even without physical aiming landmarks, participants immediately eliminated error at the primary target after being instructed to re-aim (Fig. 8H, middle, yellow). Remarkably however, without the physical aiming target, these participants did not exhibit an implicit drift in reach angle at any point during the adaptation period, and exhibited only a small implicit aftereffect during the washout period (Fig. 8H, right, t(9)=3.11, p=0.012, Cohen’s d=0.985). In fact, the aftereffect was approximately 3 times larger when participants aimed towards a physical target during adaptation than when this target was absent (Fig. 8H, right, aftereffect; two-sample t-test, t(18)=2.85, p=0.012, Cohen’s d=0.935).

Thus, these data suggested a remarkable depth to the implicit system’s response to error. While implicit adaptation was greatest in response to a target error, removal of the physical target still resulted in what appeared to be SPE-driven learning, albeit to a smaller degree.

## Discussion

Sensorimotor adaptation benefits from learning in two parallel systems: one that has access to explicit knowledge^11,51^, and another that relies on implicit, unconscious correction^12,13,45^. Here we show that each system is responsive to task-related errors between the subject’s cursor and the target^17,23^. In such cases, when the error is shared competition occurs between these systems, such that when the explicit system increases its response, errors are more rapidly depleted, thus decreasing the driving force for implicit adaptation as in Fig. 1A. This model suggests that an explicit strategy can potentially mask changes in implicit error sensitivity (Fig. 4). Indeed, suppressing the explicit strategy unveiled strong increases (Fig. 6) and decreases (Fig. 7) in putative implicit adaptation that were consistent with two hallmarks of learning: savings and interference.

However, in various cases, this task error could not explain implicit adaptation by itself. For example, when participants aimed their hand to one visual target, but the cursor to another visual target, the implicit system appeared to balance two errors (Fig. 8): an error with respect to the primary target, and an error with respect to the aimed target, an SPE. These two errors were coupled together such that decreases in error with respect to the aimed target would increase error with respect to the primary target. Thus, the data suggested a second way that the implicit system can exhibit competition: two separate implicit learning modules can interfere with one another when they try to solve conflicting objectives (Fig. 1B).

Describing sensorimotor adaptation in terms of explicit and implicit contributions is important because these systems may rely on different neural structures. Explicit learning mechanism are likely dependent on cortical involvement^43,52,53^, whereas implicit learning mechanisms at least partly engage the cerebellum^7,20,54–58^. Our results suggest that in some learning contexts, these two systems can compete with each other, as they strive to respond to a common error.

### Flexibility in the implicit response to error and the properties of savings and interference

When two similar sensorimotor perturbations are experienced in sequence, the rate of relearning is enhanced during the second exposure^28,29,32,49,59^. This hallmark of memory^60,61^ is referred to as savings. Savings is often quantified based on differences in the learning curves for each exposure^28,34^, or the rate of adaptation^62^. While these conventions are intuitive, they are based on an important underlying assumption: when one learning component’s properties change, its contribution to overall adaptation will also change. Here we describe why this intuition may not always be true.

The state space model^9,39,40^ quantifies behavior using two process: learning and forgetting. This model describes savings as a change in one’s sensitivity to error^29,32,48^. When similar errors are experienced on consecutive trials, the brain becomes more sensitive to their occurrence and responds more strongly on subsequent trials^37,48,63^. Generally, as error sensitivity increases, so too does the rate at which we adapt to the perturbation (e.g., High PT trials in Fig. 4). However, under certain circumstances, changes in one’s implicit sensitivity to error may not lead to differences in measured behavior (e.g., Low PT trials in Fig. 4).

The reason is competition. If implicit systems adapt to target errors (Eq. (1)), they are altered not solely by the rotation but also explicit strategy. When strategy is enhanced, it reduces the error available for implicit learning. Therefore, although the implicit system may become more sensitive to error, this increase in sensitivity is canceled out by the decrease in error size. If true, this would mean that implicit processes can change in ways that are hidden within measured behavior.

For example, recent lines of evidence have suggested that increases in learning rate depend solely on the explicit recall of past actions. Implicit adaptation does not seem to contribute to faster re-learning, whether its magnitude is measured through verbal reports^34^, or by restricting movement preparation time^28,33^ (Fig. 4). These data might suggest that the implicit system is unaltered by past experience. However, when reaction time is limited during both exposures, thus suppressing explicit contributions to behavior, we found that the implicit system exhibited savings (Fig. 6). This would be consistent with recent evidence that savings requires the presence of task-related errors^17^, which can be siphoned away by the explicit system. Thus, what appears to be a disconnect between studies that have detected increases in only the explicit learning rate^28,33–36^, and studies that have detected increases in the implicit learning rate^17,37,38^, may actually be consistently described by the competition equation (Eq. (4)).

This competition equation can be used to construct a map that describes how implicit adaptation should change based on the properties of implicit and explicit systems. When both implicit and explicit systems become more sensitive to error, the explicit response can hide changes in the implicit response (Fig. 5B, Region C). In fact, drastic enhancement in explicit adaptation could even lead to a decrease in implicit learning, even when implicit error sensitivity has increased (Fig. 5B, Region B). Indeed this prediction might explain cases whereby re-exposure to a rotation increases explicit strategies, but appears to attenuate implicit learning^33,36,64^. For example, a recent study by Huberdeau and colleagues^33^, seven exposures to a rotation led to caching of the explicit strategy, with a simultaneous decrease in the implicit aftereffect. However, such a mechanism cannot account for decreases in implicit learning seen in response to invariant error-clamp perturbations^36^, which presumably are free of explicit strategy.

Recent studies have shown that with multiple exposures to a visuomotor rotation, the explicit response to the perturbation can be cached and expressed at lower reaction times^33,47^. Could caching of an explicit strategy have contributed to the savings we measured under reaction time constraints in Experiment 3 (Fig. 6)? This possibility seems unlikely. First, there appears to be little such caching after only two exposures to a rotation. Otherwise, Haith and colleagues^28^ should also have observed savings on Low PT trials. In addition, the rotation occurred at four separate targets in Experiment 3, but only one target in Haith and colleagues. Lastly, reaction time constraints in Experiment 3 induced shorter reach latencies (nearing 200 ms), than those used by Haith and colleagues (300 ms). These conditions would be expected to suppress explicit caching. Nevertheless, future studies are needed to better understand the conditions (e.g., number of targets, reaction time constraints) that permit caching of the explicit process, and how these cached responses interact with implicit learning.

Finally, it is important to distinguish between reductions in implicit adaptation which appear to be driven by explicit suppression, versus those that are caused by a direct impairment in the implicit response to error. For example, when two opposing perturbations are experienced sequentially, the response to the second exposure is impaired by anterograde interference^9,25,27,65^. Recently, we linked these impairments in learning rate to a transient reduction in error sensitivity which recovers over time^26^. Here, we limited reaction time to isolate the potential implicit contributions to this impairment. Impairments in the implicit system were large and long-lasting (Fig. 7C), persisting even after 24 hours.

Interestingly, when we performed a similar experiment without restricting reaction time^26^, we found a smaller impairment in learning rate that almost fully recovered after 24 hours (Fig. 7C, no limit). These differences might suggest that uninhibited explicit strategies compensate for lingering deficits in implicit adaptation. In fact, Leow and colleagues^17^ recently demonstrated that prior exposure to task errors in one direction increases the rate at which participants explicitly adapt to a visuomotor rotation in the opposite direction, suggesting that explicit strategies might exhibit improvements rather than impairments during interference protocols. However, it is important to point out that our reaction time-limited experiment in Fig. 7, differed from our earlier work^26^ (see Methods; reaching versus pointing as well as differences in trial count). Thus, our data motivate the need for future experiments to understand how explicit strategies contribute to adaptation during anterograde interference.

### Competition-driven enhancement and suppression of implicit adaptation

Our data caution that when implicit learning increases or decreases, this does not necessarily mean that the implicit system has altered its response to error.

For example, when participants are made aware of a visuomotor rotation before it is introduced, their explicit response is drastically enhanced^15,16^. These increases in explicit strategy are coupled to decreases in implicit adaptation. A similar phenomenon can be observed in other experiments where participants are provided with visual landmarks scattered on either side of the target. When participants use these landmarks to report their intended aiming direction, reporting frequency increases explicit strategy use, but decreases implicit adaptation^66–68^. Furthermore, participants themselves exhibit varying degrees of strategy, leading to negative subject-to-subject associations between implicit and explicit learning^15,16,41^ (Fig. 3).

Given these changes in implicit adaptation, it may at first seem surprising that in some cases, implicit learning remains constant across large changes in perturbation magnitude^15,69^. For example, in Neville and Cressman^15^, while awareness decreased implicit adaptation, the implicit aftereffect was mostly invariant across each rotation size (Fig. 1). Notably, the competition equation (Eq. (4)) can again account for this observation. This equation shows that the driving force for adaptation is not the size of the rotation alone, but rather the difference between the rotation and explicit strategy (Fig. S1D).

This competition between implicit and explicit adaptation helps to reveal the errors which drive implicit learning. This competitive relationship (Eq. (4)) naturally arises when implicit systems are driven by errors in task outcome (Eq. (1)), but not errors between the cursor and intended aiming angle (Eq. (2)). We can observe these negative interactions not solely when enhancing explicit strategy, but also when suppressing re-aiming. For example, in cases where perturbations are introduced gradually, thus reducing conscious awareness, implicit “procedural” adaptation appears to increase^38,42,70^ (Fig. 2). Similarly, when participants are required to move with minimal preparation time, thus suppressing time-consuming explicit re-aiming^28,41,47^, the total extent of implicit adaptation also appears to increase^37,41^.

Lastly, competition may help to describe not only why implicit learning can vary across two experimental conditions, but also across individuals within a single experiment as in Fig. 3H. In one prime example, Miyamoto and colleagues^14^ exposed participants to a sum-of-sines rotation. Curiously, participants with more vigorous explicit responses to the perturbation exhibited less vigorous implicit learning. In a second example, Fernandez-Ruiz and colleagues^41^ observed that participants who increased their movement preparation time rapidly counteracted a rotation, but also exhibited smaller aftereffects during washout. And as a third example, when Bromberg et al.^68^ measured eye movements during adaptation, participants who tended to look towards their re-aiming locations not only exhibited greater explicit strategies, but less implicit adaptation.

In other words, participants that used cognitive strategies to adapt exhibited less procedural learning^41^. To explain these individual correlations, Miyamoto et al.^14^ suggested that there may be an intrinsic relationship between implicit and explicit sensitivity to error: when an individual’s explicit error sensitivity is high, their implicit error sensitivity is low. Here our results describe a different way to account for the same observation (Fig. 3H). In Experiment 1, we used the competition equation (Eq. (4)) to predict each individual’s implicit adaptation from their measured explicit strategy, assuming each participant had the same sensitivity to error. This one equation accurately accounted for the inverse relationship between implicit and explicit aftereffects. Thus, negative individual-level correlations between implicit and explicit adaptation can arise from subject-to-subject variation in strategy, even when implicit error sensitivity is invariant across participants.

Finally, it is important to consider how generalization may have altered our implicit learning measures. Earlier studies have shown that when participants are asked to report their aiming direction using a ring of visual landmarks, implicit learning generalizes around the reported aiming direction^71,72^. Thus, participants who aim further away from the target may show smaller implicit adaptation when asked to “move straight to the target” simply due to generalization. However, the expected magnitude of this effect (≈5°; see Fig. S5B for aim-target displacement^71^ of 22.5° and S3A for aim-target displacement^72^ of 30°) does not seem large enough to account for the large variation we measured in implicit adaptation (ranges of 17° in Fig. 2F, 32° in Fig. 3D, 14° in Fig. 3H, 17° in Fig. 3L). In the studies considered here, participants trained at either 3 (Fig. 1), 4 (Figs. 3H&L), 8 (Fig. 3D), or 12 (Fig. 2) targets, as opposed to 1 target in these earlier generalization studies^71,72^ (Figs. S3A and S3B). Thus, generalization-based decreases in implicit learning would likely be smaller in the current work, given that the generalization function widens with additional training targets^73,74^.

Along these lines, Neville and Cressman^15^ asked whether implicit learning varied across their 3 training targets, 2 of which corresponded with an “aim solution”, 1 of which did not; they did not find any change in implicit learning across each target. In addition to differences in training targets, the studies considered here did not use aiming reports to measure explicit learning, which were employed on each trial to measure aim direction in the earlier generalization studies. This may play another important role in the generalization function. For example, in these earlier generalization studies implicit learning measured via reporting was larger than that measured when reaching straight to the target (Fig. S5C), due to generalization. However, in Experiment 1, when we asked participants to report their aim at the end of adaptation, we found greater implicit learning on the straight-ahead reaching probes, than in the aim reports (Fig. S5E), opposite the generalization expectation. A similar phenomenon was noted recently when aim reports were used sparsely during adaptation^75^ (Fig. S5D). All in all, while it does not seem that generalization played a major role in our primary results, future studies are needed to measure how generalization may differ across tasks, as well as different types of error signals (e.g., target error vs. SPE).

### Error sources that drive implicit adaptation

Mazzoni and Krakauer^12^ exposed participants to a visuomotor rotation, but also provided instructions for how to re-aim their hand to achieve success. While participants immediately used this strategy to move the cursor through the target, the elimination of task error failed to stop implicit adaptation. These data suggested that implicit systems responded to errors in the predicted sensory consequence of their actions^20,76^, rather than errors in hitting the target.

However, such a model, where implicit systems learn solely based on the angle between aiming direction and the cursor (Eq. (2)), could not account for the implicit-explicit interactions we observed in some of the data (Figs. 1–3). These interactions could only be described by an implicit error source that is altered by explicit strategy, such as the angle between the cursor and the target (Eq. (1)). For example, in Experiments 1&2, participants did not aim straight to the target, but rather adjusted their aiming angle by 5-20° (Fig. 3). These changes in re-aiming appeared to alter implicit adaptation via errors between the cursor and the target. This target-cursor error source (Eq. (1)) used in our state-space model (Eq. (3)) appeared to provide an accurate account of short-term visuomotor adaptation across a number of studies^14–16,24,37,41,42^.

We do not mean to suggest however, that implicit adaptation is solely driven by a single target error. In fact, there are many cases where this idea fails^11,12,21^, beyond the Mazzoni and Krakauer study. We speculate that one feature which alters implicit learning is the simultaneous presence of multiple visual targets. In Figs. 1–7, there was only one visual target on the screen at a time. However, in Mazzoni and Krakauer (Fig. 8), there were two important visual targets: the adjacent target towards which participants explicitly aimed their hand, and the original target towards which the cursor should move. Thus, in theory there were two potential visual target errors. Interestingly, when we considered the possibility that the implicit system adapted to both errors at the same time, we could more completely account for these earlier data (Fig. 8F).

The idea that both kinds of visual error (cursor with respect to the primary target, and cursor with respect to the aimed target) drive implicit learning, could potentially help to describe other confounding observations. For example, in cases where landmarks are provided to report explicit aiming^11,24,72^, target-cursor error is often rapidly eliminated, but implicit adaptation continues to increase over time. Our dual-error model (Eq. (6)) would explain this continued adaptation based on persistent aim-cursor error.

However, the nature of this aim-cursor error remains rather uncertain. For example, while this error source generates strong adaptation when the aim location coincides with a physical target (Fig. 8H, instruction with target), implicit learning is observed even in the absence of a physical aiming landmark^21^ (Fig. 8H, instruction without target), albeit to a smaller degree. This latter condition strongly implicates an SPE learning mechanism. Thus, it may be that the aim-cursor error is actually an SPE that is enhanced by the presence of a physical target. In this view, implicit learning is driven by a target error module and an SPE module that is enhanced by a visual target error^17,23,77^. These various implicit learning modalities are likely strongly dependent on both implicit and explicit contexts, in ways we do not currently understand.

We speculate that the cerebellum might play an important role in this model of implicit adaptation^55,57,78–80^. Current models propose that complex spikes in Purkinje cells (P-cells) in the cerebellar cortex lead to LTD (Marr-Albus-Ito hypothesis). These complex spikes are reliably evoked by olivary input in response to a sensory error^79,81,82^. However, different P-cells are activated by different error directions, thus organizing P-cells into error-specific subpopulations^81,82^. Therefore, our model suggests that two different sources of error might simultaneously transduce learning in two different P-cell subpopulations, which then combine their adapted states into a total implicit correction at the level of the deep nuclei. Thus, errors based on the original target, and the aiming target, might simultaneously activate two implicit learning modules in the cerebellum (Fig. 8G).

Alternatively, it is equally possible that these aim-cursor errors and target-cursor errors engage separate brain regions both inside and outside the cerebellum. In this view, an interesting possibility is that patients with cerebellar disorders^20,54,56,83,84^ may have learning deficits specific to one error but not the other^56^. These possibilities remain to be tested.

## Methods

Here we describe the experiments and corresponding analysis reported in the main text. Much of this work involves reevaluation of earlier literature; this includes data from Haith and colleagues^28^ in Figs. 4&6, data from Lerner and Albert et al.^26^ in Fig. 7, data from Neville and Cressman^15^ in Fig. 1, data from Saijo and Gomi^42^ in Fig. 2, data from Fernandez-Ruiz et al.^41^ in Fig. 3, data from Mazzoni and Krakauer^12^ in Fig. 8, and data from Taylor and Ivry^21^ in Fig. 8. Furthermore, some generalization data^71,72^ was considered in Fig. S5. The relevant details of these studies are summarized in the sections below alongside the new data collected for this work (Experiments 1-4).

### Participants

A detailed description of participants in Haith and colleagues^28^ (n=14), Lerner and Albert et al.^26^ (n=34 for 5 min and 24 hr groups), Neville and Cressman^15^ (n=63), and Mazzoni and Krakauer^12^ (n=18), Saijo and Gomi^42^ (n=9 for abrupt, n=9 for gradual), Fernandez-Ruiz et al.^41^ (n=9), and Taylor and Ivry^21^ (n=10 for instruction with visual target, n=10 for instruction without visual target) are described in their respective papers. All volunteers (ages 18-62) in Experiments 1-4 were neurologically healthy and right-handed. Experiment 1 include n=9 participants (5 Male, 4 Female) in the No PT limit group and included n=13 participants (6 Male, 7 Female) in the PT Limit group. Experiment 2 included n=17 participants (10 Male, 7 Female). Experiment 3 included n=10 participants (6 Male, 4 Female). Experiment 4 included n=20 participants (10 Male, 10 Female). Experiments 1-4 were approved by the Institutional Review Board at the Johns Hopkins School of Medicine.

### Apparatus

In Experiments 1, 3, and 4 participants held the handle of a planar robotic arm and made reaching movements to different target locations in the horizontal plane. The forearm was obscured from view by an opaque screen. An overhead projector displayed a small white cursor (diameter = 3mm) on the screen that tracked the motion of the hand. Throughout testing we recorded the position of the handle at submillimeter precision with a differential encoder. Data were recorded at 200 Hz. Protocol details were similar for Haith and colleagues^28^, Neville and Cressman^15^, Saijo and Gomi^42^, and Fernandez-Ruiz et al.^41^ in that participants gripped a two-link robotic manipulandum, were prevented from viewing their arm, and received visual feedback of their hand position in the form of a visual cursor. In Lerner and Albert et al.^26^, participants performed pointing movements with their thumb and index finger while gripping a joystick with their right hand. In Mazzoni and Krakauer^12^, participants rotated their hand to displace an infrared marker placed on the index finger. In Taylor and Ivry^21^, hand position was tracked via a sensor attached to the index finger while participants made horizontal reaching movements along the surface of a table. In Experiment 2, participants were tested remotely on their personal computer. They moved a cursor on the screen by sliding their index finger along the track pad.

### Visuomotor rotation

Experiments 1-4 followed a similar protocol. At the start of each trial, the participant brought their hand to a center starting position (circle with 1 cm diameter). After maintaining the hand within the start circle, a target circle (1 cm diameter) appeared in 1 of 4 positions (0°, 90°, 180°, and 270°) at a displacement of 8 cm from the starting circle (in Experiment 2, 8 targets were actually used, spaced in increments of 45°).

Participants then performed a “shooting” movement to move their hand briskly through the target. Each experiment consisted of epochs of 4 trials (or 8 trials for Experiment 2) where each target was visited once in a pseudorandom order. Participants were provided audiovisual feedback about their movement speed and accuracy. If a movement was too fast (duration < 75ms) or too slow (duration > 325ms) the target turned red or blue, respectively. If the movement was the correct speed, but the cursor missed the target, the target turned white. Successful movements were rewarded with a point (total score displayed on-screen), an on-screen animation, and a pleasing tone (1000 Hz). If the movement was unsuccessful, no point was awarded and a negative tone was played (200 Hz). Participants were instructed to obtain as many points as possible throughout the experimental session.

Once the hand reached the target, visual feedback of the cursor was removed, and a yellow marker was frozen on-screen to provide static feedback of the final hand position. At this point, participants were instructed to move their hand back to the starting position. The cursor remained hidden until the hand was moved within 2 cm of the starting circle.

Movements were performed in one of three conditions: null trials, rotation trials, and no feedback trials. On null trials, veridical feedback of hand position was provided. On rotation trials, the on-screen cursor was rotated relative to the start position. On no feedback trials, the subject cursor was hidden during the entire trial. No feedback was given regarding movement endpoint, accuracy, or timing.

As a measure of adaptation, we analyzed the reach angle on each trial. The reach angle was measured as the angle between the hand and the target (relative to the start position), at the moment where the hand exceeded 95% of the target displacement.

Experiments in Haith and colleagues^28^, Lerner and Albert et al.^26^, Neville and Cressman^15^, Taylor and Ivry^21^, Saijo and Gomi^42^, Fernandez-Ruiz et al.^41^, and Mazzoni and Krakauer^12^ were collected using similar, but separate protocols. For a full description of these paradigms, please consult the corresponding manuscripts. Important differences between these experiments and the rotation protocol mentioned above are briefly described in the sections below.

### Statistics

Parametric (t-test) and nonparametric (Wilcoxon signed-rank test) tests were performed in MATLAB R2018a. For these tests, we report the p-value, and Cohen’s d as a measure of effect size.

### Competition Map

To illustrate the way implicit and explicit systems might interact, we used a state space model (Eqs. (1–3)) where implicit and explicit learning were driven by target errors. Similar to the implicit system described in Eq. (3), we modeled explicit learning as a process of learning and forgetting^14,24^:

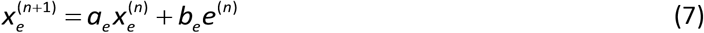

Here, *a_e_* and *b_e_* represent the explicit system’s retention factor and error sensitivity. Together Eqs. (3) and (7) describe how implicit and explicit systems adapt to error between the target and cursor (Eq. (1)).

Because implicit and explicit systems share a common error source in this target error model, their responses will exhibit competition. That is, increases in explicit adaptation will necessarily be coupled to decreases in implicit adaptation. To summarize this interaction, we created a competition map. The competition map describes common scenarios in which the goal is to compare two different learning curves. For example, one might want to compare the response to a 30° visuomotor rotation under two different experimental conditions. Another example would be savings, where we compare adaptation to the same perturbation at two different timepoints. In these cases, it is common to measure the amount of implicit and explicit adaptation, and compare these across conditions or timepoints.

The critical point is that changes in the amount of implicit adaptation reflect the modulation of both implicit and explicit responses to error. To demonstrate this idea, we needed a way to quantify the amount of implicit adaptation. For this, we chose the steady-state amount of implicit learning. As described in the main text, the steady-state level of implicit adaptation can be derived from Eqs. (1–3). This derivation resulted in the competition equation shown in Eq. (4). Note that Eq. (4) predicts the steady-state level of implicit learning from the implicit retention factor, implicit error sensitivity, mean of the perturbation, and critically, the steady-state explicit strategy. If the explicit system is also described using a state space model as in Eq. (7), it is easy to show that Eq. (4) can be equivalently expressed in terms of the implicit and explicit learning parameters according to Eq. (8):

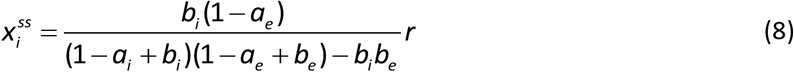

Eq. (8) provides the total amount of implicit adaptation as a function of the retention factors, *a_i_* and *a_e_*, as well as the error sensitivities, *b_i_* and *b_e_*. We used Eq. (8) to construct the competition map in Fig. 5A, by comparing the total amount of implicit learning across a reference condition and a test condition.

For our reference condition, we fit our state space model to the mean behavior in Haith et al.^28^ (Fig. 4B, Day 1, left). This model best described adaptation during the first perturbation exposure using the parameter set: *a_s_*=0.9829, *a_f_*=0.9278, *b_s_*=0.0629, *b_f_*=0.0632. Next, we imagined that implicit error sensitivity and explicit error sensitivity differed across the reference and test conditions. On the x-axis of the map, we show a percent change in *b_i_* from the reference condition to the test condition. On the y-axis of the map, we show a percent change in *b_e_* from the reference condition to the test condition. The retention factors were held constant across conditions. Then for each condition we calculated the total amount of implicit learning using Eq. (8). The color at each point in the map represents the percent change in the total amount of implicit learning from the reference condition to the test condition.

As described in the main text, the competition map (Fig. 5A) is composed of several important regions (Fig. 5B). In Region A, there is a decrease in implicit error sensitivity (from reference to test) as well as a decrease in the total amount of implicit adaptation predicted by Eq. (8). In Region B, Eq. (8) predicts a decrease in implicit adaptation, despite an increase in implicit error sensitivity. In Region D, there is an increase both in implicit error sensitivity as well as steady-state implicit learning. In Region E, there is an increase in implicit adaptation, despite a decrease in implicit error sensitivity. Finally, Region C shows cases where there are changes in implicit error sensitivity, but the total absolute change in implicit adaptation (Eq. (8)) is less than 5%. To determine this region, we solved for the linear bounds that describe a 5% increase or a 5% decrease in the output of Eq. (8).

### Neville & Cressman (2018)^15^

To understand how enhancing explicit strategy might alter implicit learning, we considered data collected by Neville and Cressman^15^. Here the authors tested how awareness of a visuomotor rotation altered the adaptation process. To do this, participants (n=63) were divided into one of many groups. In the instructed groups (Fig. 1E, yellow) the nature of the perturbation as well as a compensatory strategy was provided to the participants prior to the introduction of the perturbation. In other groups, no instruction was provided (Fig. 1E, gray). During rotation periods, participants reached to three potential targets. Implicit and explicit contributions to behavior were measured at 4 different periods using “inclusion” and “exclusion” trials. During exclusion trials, the authors instructed participants to reach (without visual feedback) as they did during the baseline period prior to perturbation onset (without using any knowledge of the perturbation gained thus far). During inclusion trials, the authors instructed participants to reach (without visual feedback) using all knowledge gained about the perturbation. In this way, the aftereffect measured on exclusion trials served as a measurement of implicit adaptation, and the difference in aftereffects measured on inclusion and exclusion trials served as a measurement of explicit adaptation.

At the start of the experiment all participants performed a baseline period without a rotation for 30 trials. Baseline implicit and explicit reach angles were then assayed using inclusion and exclusion trials. At this point, participants in the strategy group were briefed about the perturbation with an image that depicted how feedback would be rotated, and how they could compensate for it. Then all groups were exposed to the first block of a visuomotor rotation for 30 trials. Some participants experienced a 20° rotation (Fig. 1E, left), others a 40° rotation (Fig. 1E, middle), and others a 60° rotation (Fig. 1E, right). After this first block, implicit and explicit learning were assayed with inclusion and exclusion trials. This was followed by a second perturbation block, and another round of inclusion/exclusion trials. Finally, the experiment ended with a third perturbation block and a final round of inclusion/exclusion trials.

Here we focused on the measures of implicit and explicit adaptation obtained from inclusion and exclusion trials at the end of the final block. To obtain these data, we extracted the mean participant response and the associated standard error of the mean, directly from the primary figures reported by Neville and Cressman^15^ using Adobe Illustrator CS6. The implicit and explicit responses in all 6 groups are shown in Fig. S1. The marginal effects of instruction (average over rotation sizes) and rotation size (average over instruction conditions) are shown in Figs. 1F and 1G respectively.

Finally, we tested whether the competition equation (Eq. (4)) or independence equation (Eq. (5)) could account for the levels of implicit learning observed across rotation magnitude and awareness conditions. To do this, we used a bootstrapping approach. Using the mean and standard deviation obtained from the primary figures, we sampled hypothetical explicit and implicit aftereffects for 10 participants. We then calculated the mean across these 10 simulated participants. After this, we used *fmincon* in MATLAB R2018a to find an implicit error sensitivity that minimized the following cost function:

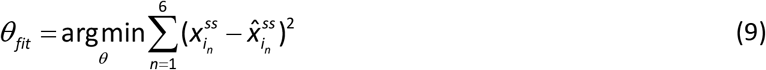

This cost function represents the difference between the simulated level of implicit adaptation, and the amount of implicit learning that would be predicted for a given perturbation size and simulated explicit adaptation, according to our competition framework (Eq. (4)) or independence framework (Eq. (5)). For this process, we set the implicit retention factor to 0.9565 (see *Measuring properties of implicit learning*). Therefore, only the implicit error sensitivity remained as a free parameter. In sum, we aimed to determine if a single implicit error sensitivity could account for the amount of adaptation across the no instruction group, instruction group, and each of the three perturbation magnitudes (20, 40, and 60°). The combination of instruction and perturbation magnitude yielded 6 groups, hence the upper limit on the sum in Eq. (9). We repeated this process for a total of 10,000 simulated groups.

In Fig. 1F, we show the marginal effect of instruction on the implicit aftereffect. This was obtained by averaging across each of the 3 rotation magnitudes shown in Fig. S1, for each model. In Fig. 1G we show the marginal effect on rotation size on the implicit aftereffect. This was obtained by averaging across the instructed and non-instructed conditions for each rotation size shown in Fig S1, for each model.

### Saijo and Gomi (2010)^42^

To understand how suppressing explicit strategy might alter implicit learning, we considered data collected by Saijo and Gomi^42^. In one of their experiments, the authors tested how perturbation onset altered the adaptation process. Subjects were divided into either an abrupt (n=9) or gradual group (n=9), and reached to 1 of 12 targets, which were ordered pseudorandomly in each cycle of 12 trials. After a baseline period of 8 cycles, a visuomotor rotation was introduced. The perturbation period lasted 32 cycles. After this, the perturbation was removed for 6 cycles of a washout condition. Participants were exposed to either an abrupt rotation where the perturbation magnitude suddenly changed from 0° to 60°, or a gradual condition where the perturbation magnitude increased over smaller increments (10° increments that lasted 3 cycles each; Fig. 2A).

Here, we considered why participants in the abrupt perturbation condition achieved greater adaptation during the rotation period (smaller error in Fig. 2C) but exhibited a smaller aftereffect when the perturbation was removed. Our theory suggested that this may be due to competition. If the gradual condition suppressed explicit awareness of the rotation^38^, then Eq. (4) would predict increases in implicit learning which were observed in the aftereffects measured during the washout period (where explicit strategies were disengaged). However, the SPE model (Eq. (5)) would predict the same amount of implicit adaptation: the same aftereffect in each condition.

To test these hypotheses, we simulated implicit adaptation using the state-space model in Eq. (3). In Fig. 2D, we used an SPE for the error term in Eq. (3). In Fig. 2E, we used the target error for the error term in Eq. (3). We imagined that the total reach angle was determined based on the sum of implicit and explicit learning. However, these authors did not directly measure explicit strategies. Fortunately, Neville and Cressman^15^ measured explicit strategies using inclusion and exclusion trials during a 60° abrupt rotation (yellow points, explicit aim in Figs. 2D&E).

We used these measurements in our abrupt simulations. Neville and Cressman observed that explicit strategies rapidly reached 35.5° and remained stable during adaptation. To approximate these data, we simulated abrupt explicit strategy using the exponential curve: *x_e_* = *35.5* − *10e*^−*2t*^ (Figs. 3D&E, explicit aim, black line). Note that the nature of this exponential curve is entirely inconsequential to our analysis, apart from its saturation level. Outside of the rotation period, we assumed explicit strategy was zero. This is consistent with data from Morehead et al.^34^ that showed almost immediate disengagement in aiming strategy during washout (Fig. S2). For the gradual condition, we assumed explicit strategy was zero throughout the entire experiment (Figs. 3D&E, explicit aim, gradual), as the participants remained largely unaware of the rotation. This seemed consistent with the data; gradual participants adapted approximately 40°, and exhibited an aftereffect of about 38°, indicating a re-aiming angle less than even 5°. Note, our primary results (Fig. 2F) were unchanged in a sensitivity test where we assumed 10° of re-aiming in the gradual group (not shown).

Thus, our simulations included two free parameters: error sensitivity (*b_i_*) and retention faction (*a_i_*) for the implicit system. In each simulation, we assumed that these parameters were identical across the gradual and abrupt groups. To fit these parameters, we minimized the following cost function:

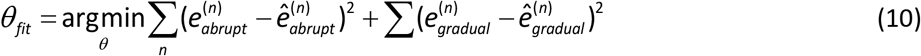

Eq. (10) is the sum of squared errors between the directional errors predicted by the model (Figs. 2D&E, directional error) and observed in the data (Fig. 2C) across all trials in the abrupt and gradual conditions. Note that each simulation incorporated variability. We simulated noisy directional errors using the standard errors shown in the data in Fig. 2C. In the explicit state, we added variability to each trial using the standard error in explicit strategy reported by Neville and Cressman^15^. For the implicit state, we used 20% of the explicit variability, given that aiming strategies are more variable than implicit corrections^14^. We repeated these simulations 20,000 times, each time resampling our noise sources and then fitting our parameter set (*a_i_* and *b_i_*) by minimizing Eq. (10) with *fmincon* in MATLAB R2018a. The mean implicit curve for the SPE learning model and target error learning model are shown in Figs. 2D and 2E respectively (implicit angle; mean ± SD). Critically, in each simulation we measured the aftereffect that occurred on the first cycle of the washout period (Figs. 2D&E, aftereffect). The mean and standard deviation in these aftereffects is reported in Fig. 2F.

Finally, note that we obtained the directional errors in Fig. 2C used in our simulations, directly from the primary figure in the original manuscript (using the GRABIT routine in MATLAB R2018a). Please also note in the actual experiment, on some trials (7.1% of all trials), the perturbation was introduced midway during the reach to test feedback corrections at only 1 target location (the 0° target). These trials were not relevant for our current analysis. Otherwise, the visuomotor rotation was applied during the entire movement as in the standard paradigm. Also note that because the authors were also analyzing feedback responses, participants made 15 cm movements, with a 0.6 second movement duration at baseline. Here, we only wanted to consider the feedforward adaptive component. Fortunately, the authors reported initial movement errors 100 ms following movement onset that could not have been altered by feedback. Therefore, we used these early measures of adaptation in the current study.

### Fernandez-Ruiz et al. (2011)^41^

In Figs. 3A-D, we show data collected and originally reported by Fernandez-Ruiz and colleagues^41^. In this experiment, participants made 10 cm reaching movements to 1 of 8 targets, pseudorandomly arranged in cycles of 8 trials. Here we report data from the unconstrained RT group described in the original manuscript. The experiment started with 3 cycles of null rotation trials, followed by 40 cycles of a 60° rotation. The experiment ended with a 20-cycle washout period (no rotation) where aftereffects were assessed. In Figs. 3B&C we show data from 2 example participants reported in the original manuscript. In Fig. 2D, the change in preparation time was calculated on the last cycle of the rotation period (relative to the baseline period). The aftereffect is the reach angle on the first cycle of the washout period. In Fig. S3, we report data from Fig. 3 of the original manuscript. Here the authors calculated the directional error and the change in preparation time across 5-cycle periods spanning the entire rotation. The points in Fig. S3 show individual subjects for the first 5 and last 5 rotation cycles. All lines show the linear regression across individual subjects in each color-coded period. Note that each line has a negative slope, indicating that participants who increased their reaction time more consistently exhibited smaller directional errors through the entire rotation period. These data were extracted directly from the primary figures reported by Fernandez-Ruiz and colleagues^41^ using Adobe Illustrator CS6. The R^2^ value reported in Fig. 2D was calculated from these extracted data.

### Experiment 1

To test whether changes in explicit strategy altered implicit learning, we recruited participants for two experiments. In the first experiment, participants adapted to a visuomotor rotation without any limits applied to preparation time (No PT limit), thus allowing participants to use explicit strategy. In a second experiment, we strictly limited preparation time in order to suppress explicit strategy (Limit PT).

Participants in the No PT limit condition began with 10 epochs of null trials (1 epoch = 4 trials), followed by a rotation period of 60 epochs. Other details concerning the experiment paradigm are described in *Visuomotor rotation*. At the end of the perturbation period, we measured the amount of implicit and explicit learning. To do this, participants were instructed to forget about the cursor and instead move their hand through the target without applying any strategy to compensate for the perturbation. Furthermore, visual feedback was completely removed during these trials. All 4 targets were tested in a randomized sequence. To quantify the total amount of implicit learning, we averaged the reach angle across all targets (Figs. 3F&H). To calculate the amount of explicit adaptation, we subtracted this measure of implicit learning from the mean reach angle measured over the last 10 epochs of the perturbation prior to the verbal instruction.

In the Limit PT group, we suppressed explicit adaptation for the duration of the experiment by limiting the time participants had to prepare their movements. To enforce this, we limited the amount of time available for the participants to start their movement after the target location was shown. This upper bound on reaction time was set to 225 ms (taking into account average screen delay). If the reaction time of the participant exceeded the desired upper bound, the participant was punished with a screen timeout after providing feedback of the movement endpoint. In addition, a low unpleasant tone (200 Hz) was played. This condition was effective in limiting reaction time (Fig. 3G, middle), even lower than the 300 ms threshold used by Haith and colleagues^28^. This experiment started with 10 epochs (1 epoch = 4 trials) of null trials. After this, the visuomotor rotation was introduced for 60 epochs. At the end of the perturbation period, we measured retention of the visuomotor memory in a series of 15 epochs of no feedback trials (Fig. 3F, no feedback).

Our goal was to test whether the putative implicit learning properties measured in the Limit PT group could be used to predict the subject-to-subject relationship between implicit and explicit adaptation in the No PT limit group (according to Eq. (4)). To do this, we measured each participant’s implicit retention factor and error sensitivity in the Limit PT condition (see *Measuring properties of implicit learning* below). We then averaged each parameter across participants. Next, we inserted these mean parameters into Eq. (4). With these variables specified, Eq. (4) predicted a specific linear relationship between implicit and explicit learning (Fig. 3H, model). We overlaid this prediction on the actual amounts of implicit and explicit adaptation measured in each No PT limit participant (Fig. 3H, black dots). We performed a linear regression across these measured data (Fig. 3H, black line, measured). We report the slope and intercept of this regression as well as the corresponding 95% confidence intervals.

The individual differences between implicit and explicit learning in Experiment 1 (Fig. 3H) could have been due uncertainty in our empirical probe (move hand through the target without re-aiming). That is, some participants may not have understood the instruction to move their hand through the target, and instead continued to aim. These participants would appear to have very little explicit strategy, and high amounts of implicit learning. Therefore, to verify our explicit measures, we considered two additional explicit markers: movement preparation time and reported strategies. In Fig. S4B, we compared explicit re-aiming with movement preparation time. That is, we calculated how much participant changed their movement preparation time after the perturbation turned on (the mean preparation time over 20 cycles following rotation onset, relative to the mean preparation time over the 3 cycles preceding rotation onset). Changes in preparation time are known to correlate with strategic re-aim^41,47^.

Lastly, we also asked participants to verbally report their explicit strategy. After the implicit probe trials, we showed each target once again, with a ring of small white landmarks placed at an equal radial distance around the screen^24^. A total of 108 landmarks was used to uniformly cover the circle. Each landmark was labeled with an alphanumeric string. Subjects were asked to report the nearest landmark that they were aiming towards at the end of the experiment in order to move the cursor through the target when the rotation was on. The mean angle reported across all 4 targets was calculated to provide an additional assay of explicit adaptation (Fig. S4A, explicit report angle). Explicit re-aiming is prone to erroneous selections where the hand is mentally rotated in the wrong direction^47^ (errors of same magnitude, opposite sign) Therefore, for individual targets where the participant reported an explicit angle in the opposite direction, we used its absolute value when calculating their explicit recalibration. These strategy report trials were used to calculate the implicit learning estimate shown in Fig. S5E.

### Experiment 2

Here, we remotely tested a very similar paradigm to the No PT limit condition in Experiment 1. Participants controlled a cursor by moving their index finger across the track pad of their personal computer. The experiment was coded in Java. To familiarize themselves with the task, participants watched a 3-minute instructional video. In this video, the trial structure, point system, and feedback structure were described. After this video, there was a practice period. During the practice period, the software tracked the participant’s reach angle on each trial. If the participant achieved success on fewer than 65% of trials (measured based on an angular target-cursor discrepancy ≤ 30°, reaction time ≤ 1 sec, and movement duration ≤ 0.6 sec), they had to re-watch the instructional video and re-do the practice period.

After the practice period ended, the testing period began. This testing period was almost identical to the No PT limit condition in Experiment 1. On each trial, participants reached to 1 of 4 targets (up, down, left, and right). Each target was visited once pseudorandomly in a cycle of 4 targets. After an initial 10-cycle null period, a 30° visuomotor rotation was imposed that lasted for 60 epochs. At the end of the rotation period, we measured implicit and explicit adaptation. The experiment briefly paused, and an audiovisual recording was played that instructed participants to not use any strategy and to move their hand straight through the target. After this, the experiment resumed, feedback was removed, and participants performed 20 cycles of no-aiming, no-feedback probe trials (Fig. 3J, no aiming).

We measured subject-to-subject correlations between implicit and explicit adaptation. For this, we calculated two implicit learning measures. The early implicit aftereffect was simply the aftereffect observed on the first no-aiming, no-feedback probe cycle (Fig. 3L). The late implicit aftereffect was the average aftereffect observed on the last 15 cycles of this no-aiming, no-feedback period (Fig. 3K). To measure explicit learning, we calculated the difference between the total amount of adaptation (mean reach angle over last 10 cycles of the rotation period) and the first cycle of the no-aiming, no-feedback period. We investigated the relationship between explicit adaptation and the early and late implicit aftereffects via linear regression in Figs. 3L and 3K respectively. For the early implicit aftereffect, we measured the 95% CI for the slope and intercept. Critically, this interval did not contain 1, indicating that the subject-to-subject correlations cannot be described by the trivial case where all participants had adapted the same amount by the end of the adaptation period (see main text).

### Haith et al. (2015)^28^

To understand how implicit and explicit processes contribute to savings, Haith and colleagues^28^ designed a forced preparation time task. Briefly, participants (n=14) performed reaching movements to two targets, T1 and T2, under a controlled preparation time scenario. To control movement preparation time, four audio tones were played (at 500 ms intervals) and participants were instructed to reach coincident with the 4th tone. On high preparation time trials (High PT), the intended target was displayed during the entire tone sequence. On low preparation time trials (Low PT), the intended target was switched approximately 300 ms prior to the 4th tone. High PT trials were more probable (80%) than Low PT trials (20%).

After a baseline period (100 trials for each target), a 30° visuomotor rotation was introduced for target T1 only. After 100 rotations trials (Exposure 1), the rotation was turned off for 20 trials. After a 24 hr break, participants then returned to the lab. On Day 2, participants performed 10 additional reaching movements without a perturbation, followed by a second 30° rotation (Target T1 only) of 100 trials (Exposure 2). The experiment then ended with a washout period of 100 trials for each target.

We quantified the amount of savings expressed upon re-exposure to the perturbation, on High PT and Low PT trials. We measured savings using two metrics. First, we measured the rate of learning during each exposure to the perturbation using an exponential fit. We fit a two-parameter exponential function to both Low PT and High PT trials during the first and second exposure (we constrained the third parameter to enforce that the exponential begin at each participant’s measured baseline reach angle). We compared the exponential learning rate using a paired t-test (Fig. 4B, 3rd column).

We also quantified savings in a manner similar to that reported by Haith and colleagues^28^; we calculated the difference between the reach angles before and after the introduction of the perturbation, during each exposure (Fig. 4C, 1st and 2nd columns). For High PT trials, we then computed the mean reach difference over the 3 trials preceding, and 3 trials following perturbation onset. Given their reduced frequency, for Low PT trials, we focused solely on the trial before and trial after perturbation onset. To detect savings, we compared the pre-perturbation and post-perturbation differences using a paired t-test (Fig. 4C, 3rd column).

Finally, we also used a state-space model of learning to measure properties of implicit and explicit learning during each exposure. We modeled implicit learning according to Eq. (3) and explicit learning according to Eq. (7). In one model fitting procedure, we modeled error according to Eq. (1) for the competitive framework. These results are shown in Fig. 4D. In a second model fitting procedure, we modeled error according to Eq. (2) for the independent framework. These results are not shown in the Fig. 4, but relevant statistical outcomes are reported in the main text.

In the model, behavior is described as the summation of implicit and explicit learning. Each system possessed a retention factor and error sensitivity. Here, we asked how implicit and explicit error sensitivity might have changed from Exposure 1 to Exposure 2. Therefore, we assumed that the implicit and explicit retention factors were constant across perturbations, but allowed a separate implicit and explicit error sensitivity during Exposures 1 and 2. Therefore, our modeling approach included six free parameters. We fit this model to the measured behavior by minimizing the following cost function using *fmincon* in MATLAB R2018a:

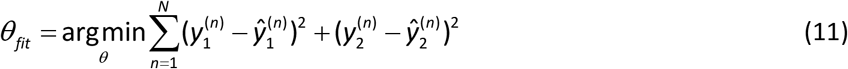

Here *y*_1_ and *y*_2_ represent the reach angles during the first and second exposure. These reach angles are composed of High PT and Low PT trials. On Low PT trials, the reach angle is equal to the implicit adaptative process. On High PT trials, the reach angle is equal to the sum of the implicit adaptive process and the explicit adaptive process.

We fit this model to individual participant behavior, in the case where implicit learning was driven by target errors (Eq. (1)), and also in the alternate case where it was driven by aim-cursor errors (Eq. (2)). We report the implicit and explicit error sensitivities for the target-error learning case in Fig. 4D, right. For this model, the predicted behavior is shown in the first two columns of Fig. 4D. We also fit the target-error (Eq. (1)) model to the mean behavior across all participants in Exposure 1 and Exposure 2. We obtained the following parameter set: *a_s_*=0.9829, *a_f_*=0.9278, *b_s,1_*=0.0629, *b_s,2_*=0.089, *b_f,1_*=0.0632, *b_f,2_*=0.1078. Note that the subscripts 1 and 2 denote error sensitivity during Exposure 1 and 2, respectively. These parameters were used for our simulations in Fig. 5 (see *Competition Map*).

### Experiment 3

In Haith et al. (2015)^28^, no savings was observed on trials where preparation time was limited (Low PT trials), consistent with the possibility that implicit learning processes are not modulated by past experiences. Here, we questioned if savings in implicit learning processes might have been suppressed by competition with explicit learning processes (see *Competition Map*). That is, if implicit and explicit processes share error sources, changes in explicit learning could mask changes in implicit learning. The way to test this possibility would be to eliminate explicit learning on all trials, to ensure that the error on each trial is expressly available for the implicit learning system. Experiment 3 tested this possibility using a limited preparation time condition.

Limiting reaction time is known to suppress explicit strategy^17,41,47^. To limit reaction time, we used the same procedure described above for Experiment 2. This condition was effective in limiting reaction time (Fig. 6B, top row), even lower than the 300 ms threshold used by Haith and colleagues^28^.

Experiment 3 used the 4-target protocol reported in *Visuomotor rotation*. Apart from that, its trial structure was similar to that of Haith et al.^28^. After a familiarization period, subjects completed a baseline period of 10 epochs (1 epoch = 4 trials for each target). At that point, we imposed a 30° visuomotor rotation for 60 epochs (Exposure 1). At the end of this first exposure, participants completed a washout period with no perturbation that lasted for 70 epochs. At the end of the washout period, subjects were once again exposed to a 30° visuomotor rotation for 60 epochs (Exposure 2).

We quantified savings in a manner consistent with Haith et al.^28^. First, we fit a two-parameter exponential function to the reach angle during Exposures 1 and 2 (third parameter was used to constrain the fit so exponential curve started at the reach angle measured prior to perturbation onset). We analyzed any change in the rate parameter of the exponential using a paired t-test (Fig. 6C, top). Second, we also tested for differences in the initial amount of learning. To do this, we calculated the difference between reach angle during Exposures 1 and 2 (Figs. 6A&B, bottom row). We then calculated the difference in reach angle (Exposure 2 - Exposure 1) during the 4 epochs preceding and 4 epochs following rotation onset. We compared these differences for Exposures 1 and 2 using a paired t-test (Fig. 6C, bottom).

### Experiment 4

Lerner and Albert et al.^26^ demonstrated that anterograde interference slows the rate of learning after 5 min (also 1 hr), but dissipates over time and is nearly gone after 24 hr. Here we wondered if this reduction in learning rate could at least be in part driven by impairments in implicit learning. Because Lerner and Albert et al.^26^ did not constrain preparation time, one would expect that participants used both implicit and explicit learning processes. In Experiment 2, we isolated the implicit component of adaptation by limiting reaction time. We used the same technique to limit reaction time reported for Experiment 2. The experiment paradigm is described in *Visuomotor rotation* above. With that said, we used 8 adaptation targets as opposed to 4 targets, to match the protocol used by Lerner and Albert et al.^26^.

The perturbation schedule is shown in Figs. 7A&B at top. We recruited two groups of participants, a 5 min group (n=9), and a 24 hr group (n=11). After familiarization, all participants were exposed to a baseline period of null trials lasting 5 epochs (1 epoch = 8 trials). Next participants were exposed to a 30° visuomotor rotation for 80 cycles (Exposure A). At this point, the experiment ended. After a break, participants returned to the task. For the 5 min group, the second session occurred on the same day. For the 24 hr group, participants returned the following day for the second session. At the start of the second session, participants were exposed to a 30° visuomotor rotation (Exposure B) whose orientation was opposite to that of Exposure A. This rotation lasted for 80 epochs.

We analyzed the rate of learning by fitting a two-parameter exponential function to the learning curve during Exposures A and B (the third parameter was used to constrain the exponential curve to start from the behavior on the first epoch of the rotation). For each participant we computed an interference metric by dividing the exponential rate of learning during Exposure B, by that measured during Exposure A (Fig. 7C, at right, blue). In addition, we also analyzed the reaction time of the participants during Exposure B (Figs. 7A&B, middle, blue).

### Lerner and Albert et al. (2020)^26^

Recently, Lerner and Albert et al.^26^ demonstrated that slowing of learning in anterograde interference paradigms is caused by reductions in sensitivity to error. Here, we re-analyze some of these data.

Lerner and Albert et al.^26^ studied how learning one visuomotor rotation altered adaptation to an opposing rotation when these exposures were separated by time periods ranging from 5 min to 24 hr. Here we focused solely on the 5 min group (n=16) and the 24 hr group (n=18). A full methodological description of this experiment is provided in the earlier manuscript. Briefly, participants gripped a joystick with the thumb and index finger which controlled an on-screen cursor. Their arm was obscured from view using a screen. Targets were presented in 8 different positions equally spaced at 45° intervals around a computer monitor. Each of these 8 targets was visited once (random order) in epochs of 8 trials. On each trial, participants were instructed to shoot the cursor through the target.

All experiment groups started with a null period of 11 epochs (1 epochs = 8 trials). This was followed by a 30° visuomotor rotation for 66 epochs (Exposure A). At this point, the experiment ended.

After a break, participants returned to the task. For the 5 min group, the second session occurred on the same day. For the 24 hr group, participants returned the following day for the second session. At the start of the second session, participants were immediately exposed to a 30° visuomotor rotation (Exposure B) whose orientation was opposite to that of Exposure A. This rotation lasted for 66 epochs. Short set breaks were taken every 11 epochs during Exposures A and B.

Here as in the earlier work^26^, we analyzed the rate of learning by fitting a two-parameter exponential function to the learning curve during Exposures A and B (the third parameter was used to constrain the exponential curve to start from the behavior on the first epoch of the rotation). For each participant we computed an interference metric by dividing the exponential rate of learning during Exposure B, by that measured during Exposure A (Fig. 7C, green). In addition, we also analyzed the reaction time of the participants during Exposure B. The mean reaction time over the first perturbation block is shown in Figs. 7A&B (middle, green traces).

### Mazzoni and Krakauer (2006)^12^

In this study, subjects sat in chair with their arm supported on a tripod. An infrared marker was attached to a ring placed on the participant’s index finger. The hand was held closed with surgical tape. Participants moved an on-screen cursor by rotating their hand around their wrist. These rotations were tracked with the infrared marker. On each trial, participants were instructed to make straight out-and-back movements of a cursor through 1 of 8 targets, spaced evenly in 45° intervals. A 2.2 cm marker translation was required to reach each target. Note that all 8 targets remained visible throughout the task.

Two groups of participants were tested with a 45° visuomotor rotation. In the no-strategy group, participants adapted as per usual, without any instructions. After an initial null period, the rotation was turned on (Fig. 8A, blue, adaptation). After about 60 cycles of adaptation, the rotation was turned off and participants performed another 60 of washout trials (Fig. 8A, blue, washout). The break between the adaptation and washout periods in Fig. 8A, no-strategy, is simply for alignment purposes.

The strategy group followed a different protocol. After the null period, participants reached for 2 movements under the rotation (Fig. 8A, 2 cycles no instruction, red). At this point, the subjects were told that they made 2 errors, and that they could counter the error by reaching to the neighboring clockwise target (all targets always remained onscreen). After the instruction, participants immediately reduced their error to zero (point labeled instruction in red, Fig. 8A). They continued to aim to the neighboring target under the rotation throughout the adaptation period. Note that the direction errors became negative. This convention indicates overcompensation for the rotation, i.e., that participants are altering their hand angle by more than their strategy aim of 45°. Towards the end of the adaptation period, participants were told to stop re-aiming, and direct their movement back to the original target (Fig. 8A, do not aim, rotation on). Then after several movements, the rotation was turned off as participants continued to aim for the original target during the washout period.

In Fig. 8A we show the error between the primary target (target 1) and cursor during the entire experiment. In Fig. 8B we show the error between the aimed target (target 2) and cursor during the adaptation period. Note that the aimed and primary targets are generally related by 45° when the strategy group is re-aiming. We observed that initial adaptation rates (over first 24 movements, gray area in Fig. 8B) were similar, but the no-strategy group ultimately achieved greater implicit adaptation. These data were all obtained by using the GRABT routine in MATLAB 2018a to extract the mean (and standard error of the mean) performance in each group from the figures shown in the primary article.

To account for behaviors, we fit 1 of 3 models to the direction error during the adaptation period shown in Fig. 8B. In all cases we modeled explicit re-aiming in the strategy group as an an aim sequence that started at zero during the initial two movements, and then 45° for the rest of the adaptation period (i.e., after the instruction to re-aim). In the no-strategy group, we modeled explicit learning as an aim sequence that remained at zero throughout the adaptation period.

In Fig. 8D, we modeled implicit learning based on the state-space model in Eq. (3) and target error term defined in Eq. (1). This target error was defined as the difference between the primary target (i.e., the initial target displayed associated with task outcome) and the cursor. In Fig. 8E, we modeled implicit learning based on the state-space model in Eq. (3) and the aim-cursor error defined in Eq. (2). This aim-cursor error was defined as the difference between the aimed target (either 0° or 45°) and the cursor. Fig. 8F, shows our third and final model. In this model, implicit learning in the strategy group was modeled using the dual-error system shown in Eq. (6). That is, there were two implicit modules, one which responded to the target errors as in Fig. 8D, and the other which responded to aim-cursor errors as in Fig. 8E. The evolution of these errors is shown in Fig. 8G. In the no-strategy group, we modeled implicit learning based on the primary target error alone and cursor.

Each model in Figs. 8D-F were fit in an identical manner. We fit the implicit retention factor and implicit error sensitivity to minimized squared error according to:

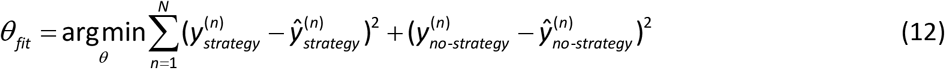

In other words, we minimized the sum of squared error between our model fit and the observed behavior across both the strategy and no-strategy groups in Fig. 8B. In other words, we constrained that each group had the same implicit learning parameters. In the case of our dual-error model in Fig. 8F, we assumed that each implicit module also possessed the same retention and error sensitivity. In sum, all model fits had two free parameters (error sensitivity and retention) which were assumed to be identical independent of instruction. This fit was performed using *fmincon* in MATLAB R2018a. The predicted behavior is shown in Figs. 8D-F at bottom. For our best model (Fig. 8F), the model behavior is also overlaid in Fig. 8B.

### Taylor and Ivry (2011)^21^

In Fig. 8H, we show data collected and originally reported by Taylor and Ivry^21^. In this experiment, participants moved their arm at least 10 cm towards 1 of 8 targets, that were pseudorandomly arranged in cycles of 8 trials. Only endpoint feedback of the cursor position was provided. The hand was slid along the surface of a table while the position of the index finger was tracked with a sensor. After an initial familiarization block (5 cycles), participants were trained how to explicitly rotate their reach angle clockwise by 45°. That is, on each trial they were shown veridical feedback of their hand position, but were told to reach to a neighboring target, that was 45° away from the primary illuminated target. After this training and another null period, the adaptation period started where the cursor position was rotated by 45° in the counterclockwise direction for 40 cycles. The first 2 movements in the rotation exhibited large errors (Fig. 8H, 2 movements no instruction). As in Mazzoni and Krakauer^12^, the participants were then instructed that they could minimize their error by adopting the aiming strategy they learned at the start of the experiment. Using this strategy, participants immediately reduced their direction error to zero.

Here we report data from two critical groups in this experiment. In the “instruction with target” group (Fig. 8H, black, n=10) participants were shown the neighboring targets during the adaptation period to assist their re-aiming. However, in the “instruction without target” group (Fig. 8H, yellow, n=10) participants were only shown the primary target; the neighboring targets did not appear on the screen to help guide re-aiming. Only participants in the “instruction with target” group exhibited the drift reported by Mazzoni and Krakauer^12^. However, both groups exhibited an implicit aftereffect (Fig. 8H, aftereffect; first cycle of washout period as reported in Fig. 4C of the original manuscript^21^).

These data were extracted directly from the primary figures reported by Taylor and Ivry^21^ using Adobe Illustrator CS6. We used the means and standard deviations for our statistical tests on the implicit aftereffect in Fig. 8H.

### Generalization studies

In our Discussion, we describe how generalization can alter measurements of implicit adaptation. Here we report data from many earlier studies. In Fig. S5A, we show data collected by Day et al.^72^, reported in Fig. 2 of the original manuscript. Here, participants were exposed to a 45° rotation while reaching to a single target. On each trial they were asked to report their aiming direction, using a ring of visual landmarks. In the “target” group in Fig. S5A, implicit aftereffects were periodically probed at the trained target location, by asking participants to reach to the target without aiming. In the “aim” group in Fig. S5A, implicit aftereffects were periodically probed at a target location 30° away from the trained target, consistent with the direction of the most frequently reported aim. In Fig. S5A, we show the implicit aftereffect measured on the first aftereffect trial at the end of the experiment. In Fig. S5C we again show the implicit aftereffect measured at the trained target location in the “probe” condition. The “report” condition shows the amount of implicit learning estimated by subtracting the reported explicit strategy from the reported reach angle on the last cycle of the rotation.

In Fig. S5B, we show data collected by McDougle et al.^71^, reported in Fig. 3A of the original manuscript. Here participants were also exposed to a 45° rotation while reaching to a single target. At the end of the experiment, participants were exposed to an aftereffect block where they reached 3 times to 16 different targets spaced in varying increments around the unit circle. In this aftereffect block feedback was removed and participants were told to move straight to the target without re-aiming. This aftereffect block was used to construct a generalization curve. In Fig. S5B we show data only from 2 relevant locations on this curve. The “target” condition represents aftereffects probed at the training target. The “aim” condition shows the aftereffect measured at 22.5° away from the primary target, which was the target closest to the mean reported explicit re-aiming strategy of 26.2°.

Lastly, in Fig. S5D we show data collected by Maresch et al.^75^, reported in Fig. 4b of the original manuscript. This study was informative to our discussion because they report implicit aftereffects measured using both exclusion trials (as in most of the data described in this manuscript) as well implicit aftereffects measured using aim reports. In Fig. S5D we specifically show data from the IR-E group in the original manuscript. We selected this group, because aim was only intermittently reported (4 trials for every 80 normal adaptation trials), and also because there were many adaptation targets (8 total). Thus, in most cases, participants only had to attend to a single target when reaching as in our primary results. The “probe” condition in Fig. S5D corresponds to the total implicit learning measured at the end of adaptation by telling participants to reach without re-aiming. The “report” condition in Fig. S5D corresponds to the total implicit learning estimated at the end of adaptation by subtracting the reported aim direction from the measured reach angle.

Note that data in Figs. S5A-D were extracted directly from the primary figures reported in the original manuscripts using Adobe Illustrator CS6.

### Measuring properties of implicit learning

Many of our model’s predictions depended on estimates of implicit retention factor and error sensitivity. We obtained these using the Limit PT group in Experiment 2. To calculate the retention factor for each participant, we focused on the no feedback period at the end of Experiment 2 (Figs. 8D, no feedback). During these error-free periods trial errors were hidden, thus causing decay of the learned behavior. The rate of this decay is governed by the implicit retention factor according to:

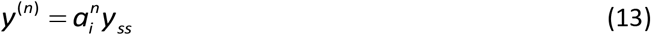

Here *y*^(n)^ refers to the reach angle on the n-th no feedback trial, and *y_ss_* corresponds to the asymptotic behavior prior to the no feedback period. We used *fmincon* in MATLAB R2018a to identify the retention factor which minimized the difference between the decay predicted by Eq. (13) and that measured during the no feedback period. We obtained an epoch-by-epoch retention factor of 0.943 ± 0.011 (mean ± SEM). Note that an epoch consisted of 4 trials, so this corresponded to a trial-by-trial retention factor of 0.985. When modeling Neville and Cressman^15^ (Fig. 1), we cubed this trial-by-trial term because each cycle consisted of 3 different targets (final retention factor of 0.9565).

Next, we measured implicit error sensitivity in the Limit PT group during rotation period trials. To measure implicit error sensitivity on each trial, we used its empirical definition:

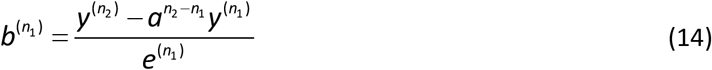

Eq. (14) determines the sensitivity to an error experienced on trial *n_1_* when the participant visited a particular target T. This error sensitivity is equal to the change in behavior between two consecutive visits to target T, on trials *n_1_* and *n*_2_ divided by the error that had been experienced on trial *n*_1_. In the numerator, we account for decay in the behavior by multiplying the behavior on trial *n*_1_ by a decay factor that accounted for the number of intervening trials between trials *n*_1_ and *n*_2_. For each target, we used the specific retention factor estimated for that target with Eq. (13).

Using this procedure, we calculated implicit error sensitivity as a function of trial in Experiment 2. To remove any potential outliers, we identified error sensitivity estimates that deviated from the population median by over 3 median absolute deviations within windows of 10 epochs. As reported by Albert and colleagues^37^, implicit error sensitivity increased over trials. Eqs. (4) and (5) require the steady-state implicit error sensitivity observed during asymptotic performance. To estimate this value, we averaged our trial-by-trial error sensitivity measurements over the last 5 epochs of the perturbation. This yielded an implicit error sensitivity of 0.346 ± 0.071 (mean ± SEM).

## Acknowledgements

This work was supported by grants from the National Institutes of Health (R01NS078311, F32NS095706), and the National Science Foundation (CNS-1714623).

---

**Figure S1.**
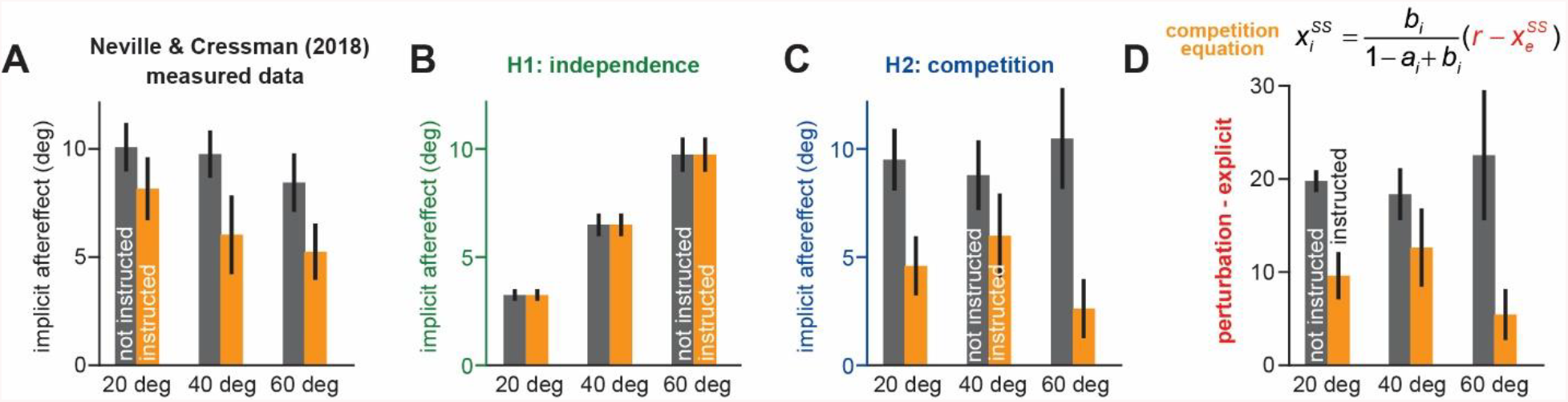
Changes in implicit adaptation in response to awareness and rotation size. Data reported from Neville and Cressman (2018)^15^. **A.** Participants were separated into 1 of 6 groups. Groups differed based on verbal instruction (instructed yellow; non-instructed gray) and rotation magnitude (20° left; 40° middle; 60° right). Here we show implicit learning measured using exclusion trials (reach without re-aiming) at the end of adaptation. **B.** Here we show implicit aftereffects predicted by a model where implicit system learns from SPE only. **C.** Here we show implicit aftereffects predicted by a model where implicit system learns from target error only. **D**. The competition model (target error learning) predicts that implicit learning will be proportional to the difference between the rotation size and the total explicit strategy. Here we show this quantity for all 6 experimental groups. Note that model predictions in **B** and **C** assume that implicit error sensitivity and retention factor are the same across all 6 experimental groups. Error bars for data show mean ± SEM. Error bars for model predictions refer to mean and standard deviation across 10,000 bootstrapped samples.

**Figure S2.**
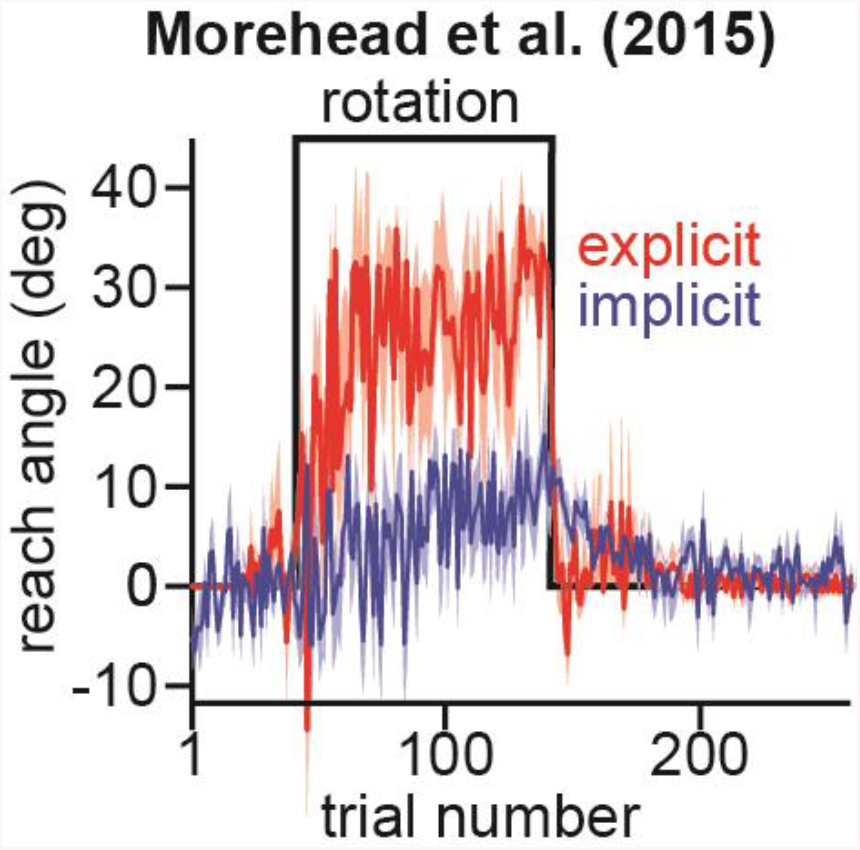
Explicit strategies are rapidly disengaged during washout. Data are reported from Morehead et al. (2015)^34^. Here participants adapted to a 45° rotation, followed by an extended washout period. Explicit learning was measured by asking subjects to report their aiming angle using a ring of visual landmarks. Implicit learning was measured as the difference between the observed reach angle and the direction of reported aim. In this task, participants reached on each trial to 1 of 4 targets. Note the sharp change in explicit angle to zero at the start of the washout period. The aftereffect during a washout period is thought to reflect implicit adaptation. This requires that explicit strategies are rapidly disengaged during washout, consistent with these data. Error bars show mean ± SEM.

**Figure S3.**
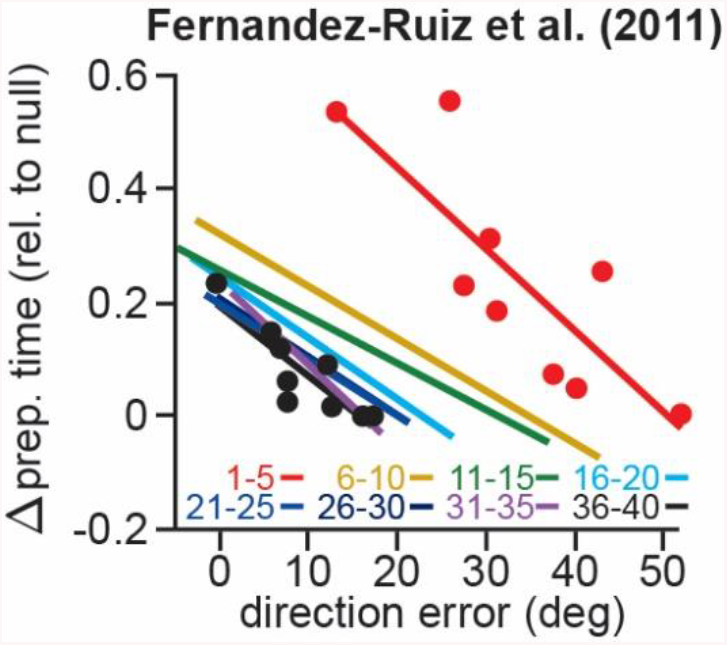
Participants that increase their preparation time exhibit greater total adaptation. Data are reported from Fernandez-Ruiz and colleagues^41^. In this experiment, participants made 10 cm reaching movements to 1 of 8 targets, pseudorandomly arranged in cycles of 8 trials. Here we report data from the unconstrained RT group described in the original manuscript. The experiment started with 3 cycles of null rotation trials, followed by 40 cycles of a 60° rotation. The authors calculated change in movement preparation time (relative to baseline period) on each trial. Here the authors calculated the directional error and the change in preparation time across 5-cycle periods spanning the entire 40-cycle rotation. The points show individual subjects for the first 5 and last 5 rotation cycles. All lines show the linear regression across individual subjects in each color-coded period. Note that each line has a negative slope, indicating that participants who increased their reaction time more consistently exhibited smaller directional errors through the entire rotation period.

**Figure S4.**
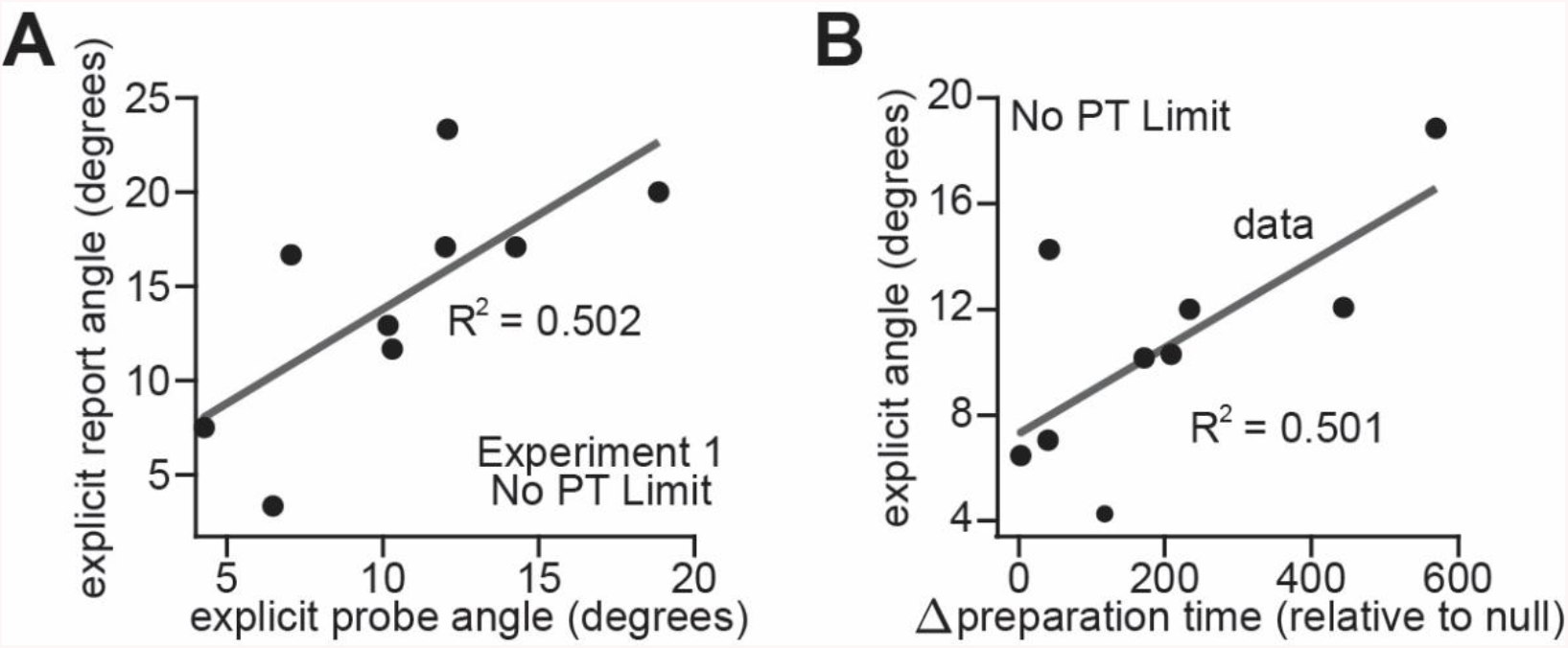
Alternate measures of explicit strategy. **A.** In the No PT limit participants in Experiment 1, we empirically measured explicit re-aiming at the end of adaptation. To do this, we instructed participants to move their hand through the target without any re-aiming. Reach angle precipitously dropped after this instruction. The total change in reach angle (averaged across all 4 targets) represented each participant’s strategic re-aiming (x-axis). To validate this empirical measure, we also asked participants to report their explicit strategies after the probe period. Participants were shown a ring of circles surrounding each target and asked to indicate which circle best represented their aiming during at the end of the experiment. This reported explicit measure averaged across all 4 targets is shown on the y-axis. Each dot represents one participant. **B.** Explicit strategies have also been shown to correlate with increases in movement preparation time. Here we show the total explicit strategy measured (via the no aiming probe trial in No PT limit in Experiment 1) as a function of change in preparation time for each individual participant. The change in preparation time was calculated as the difference between the mean preparation time over the first 20 rotation cycles and the last 3 null period cycles. The solid lines in **A** and **B** show a linear regression across individual participants.

**Figure S5.**
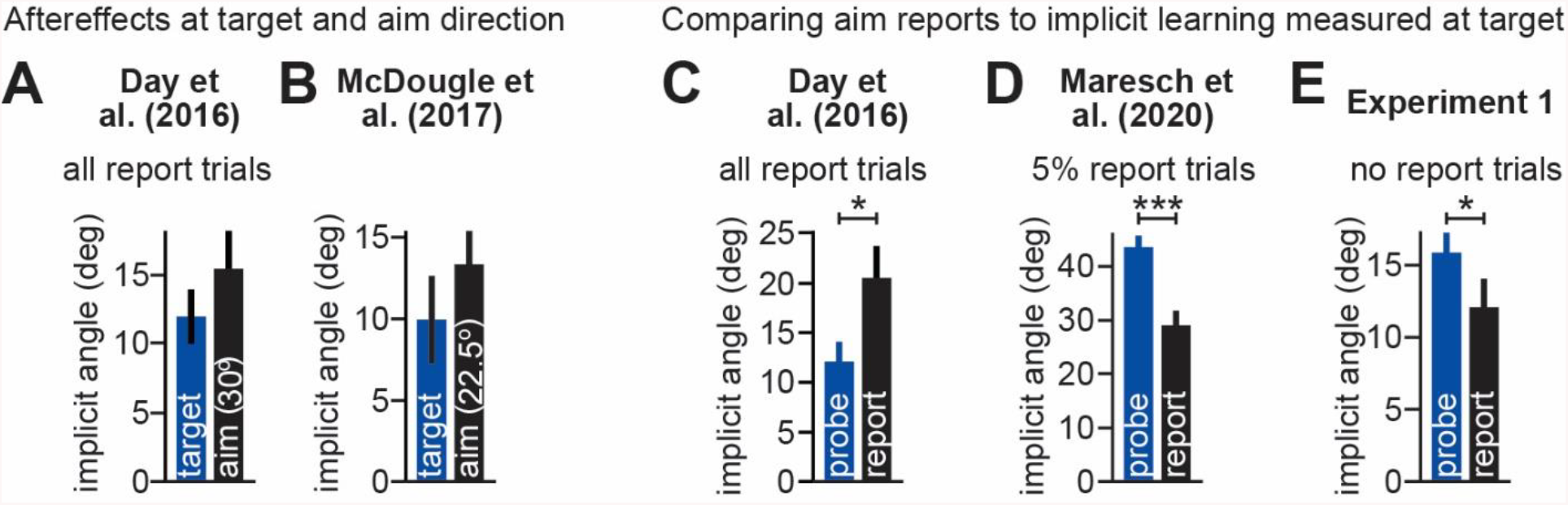
Differences in generalization across visuomotor rotation tasks. **A.** Data collected by Day et al.^72^, reported in Fig. 2 of the original manuscript. Here, participants were exposed to a 45° rotation while reaching to a single target. On each trial they were asked to report their aiming direction, using a ring of visual landmarks. In the “target” group, implicit aftereffects were measured at the trained target location. In the “aim” group, implicit aftereffects were probed at a target location 30° away from the trained target, consistent with the direction of the most frequently reported aim. Here we show data from the first aftereffect cycle after the rotation period. **B.** Similar to **A** except for data reported by McDougle et al.^71^ (Fig. 3A of the original manuscript). Participants were also exposed to a 45° rotation while reaching to a single target. At the end of the experiment, participants were exposed to an aftereffect block where participants were told to move straight to the target without re-aiming. Here we take two relevant points from the generalization curve measured at the end of learning. The “target” condition represents aftereffects probed at the training target. The “aim” condition shows the aftereffect measured at 22.5° away from the primary target, which was the target closest to the mean reported explicit re-aiming strategy of 26.2°. **C.** Data again from Day et al.^72^. The “probe” implicit learning measure is the same as **A**. The “report” condition shows the amount of implicit learning estimated by subtracting the reported explicit strategy from the reported reach angle on the last cycle of the rotation. **D.** Similar to **C**, but for the intermittent reporting (IR-E) group reported by Maresch et al.^75^ (Fig. 4b of the original manuscript). In this group aim was only intermittently reported (4 trials for every 80 normal adaptation trials). Thus, in most cases, participants only had to attend to a single target when reaching. The authors also used 8 training targets (as opposed to 1 in **A**-**C**). The “probe” condition corresponds to the total implicit learning measured at the end of adaptation by telling participants to reach without re-aiming. The “report” condition corresponds to the total implicit learning estimated at the end of adaptation by subtracting the reported aim direction from the measured reach angle. **E.** Here we report implicit learning measured using the “probe” and “report” conditions in Experiment 1, analogous to the measures described in **D**. Error bars show mean ± SEM.

